# Gapped-kmer sequence modeling robustly identifies regulatory vocabularies and distal enhancers conserved between evolutionarily distant mammals

**DOI:** 10.1101/2023.10.06.561128

**Authors:** Jin Woo Oh, Michael A. Beer

## Abstract

Gene regulatory elements drive many complex biological phenomena such as fetal development, and their mutations are linked to a multitude of common human diseases. The phenotypic impacts of regulatory variants are often tested using their conserved orthologous counterparts in model organisms such as mice. However, mapping human enhancers to conserved elements in mice remains a challenge, due to both rapid evolution of enhancers and limitations of current computational methods to detect conserved regulatory sequences. To improve upon existing computational methods and to better understand the sources of this apparent regulatory divergence, we comprehensively measured the evolutionary dynamics of distal enhancers across 45 matched human/mouse cell/tissue pairs from more than 1,000 DNase-seq experiments. Using this expansive dataset, we show that while cell-specific regulatory vocabulary is conserved, enhancers evolve more rapidly than other genomic elements such as promoters and CTCF binding sites. We observed surprisingly high levels of cell-specific variability in enhancer conservation rates, in part explainable by tissue specific transposable element activity. To improve orthologous enhancer mapping, we developed an improved genome alignment algorithm using gapped-kmer sequence features, and using the matched cell/tissue pairs, we show that this novel computational method, *gkm-align*, discovers 23,660 novel human/mouse conserved enhancers missed by standard alignment algorithms.

## Introduction

Model organisms have been indispensable for understanding the functional roles of cis-regulatory elements (CRE) in complex biological phenomena such as fetal development and pathogenesis. Some human CREs with putative functional roles have been validated by dissecting the mutational impact of their conserved orthologous counterparts in model animals^1^. However, identifying orthologous CREs, especially for distal enhancers, is computationally challenging both due to their rapid evolution and their degenerate sequence structures. Functional characterization of regulatory elements^2^ and non-coding GWAS disease associated variants^3–5^ typically begins with mapping human enhancers to mouse with sequence alignment, which suffers from low sensitivity, and thus the accurate identification of orthologous CREs has long been a bottleneck in efforts to improve our understanding of CRE function.

Enhancers are DNA sequences harboring multiple transcription factor binding sites (TFBS) and are important regulators of gene expression^6^. In spite of their importance, enhancers have evolved rapidly relative to protein-coding sequences^7^ and promoters^8–10^. Since DNA motifs for TFBS are often degenerate and spacing between TFBS typically do not contribute to enhancer function, nucleotide substitutions can accumulate more readily without significant functional changes in enhancers^11^, and this flexibility likely aids regulatory evolution. Further, duplication of redundant TFBS and accumulation of enhancer mutations can lead to turnover of TFBS through stabilizing selection^11^. This flexibility also applies on a larger scale to combinations of enhancers within intergenic loci, as enhancer function is weakly constrained by position relative to the target promoter, and enhancers are often accompanied by redundant *shadow* enhancers that regulate the same target genes^12–14^. As a result, the functional and mechanistic properties of enhancers likely have facilitated their rapid turnover throughout evolutionary history while maintaining DNA binding specificities of TFs^8,15^. Our analysis below supports this picture of rapid enhancer evolution in the context of conserved TF binding specificity across a broad range of cell types.

Previously, several groups have observed that many putative enhancers, marked by chromatin accessibility^8^, TF-binding^10^, and/or histone modifications^9^ characteristic of cis-regulatory elements (e.g., H3K27ac, H3K4me1/3), lack functional conservation at orthologous loci of distant mammals predicted by conventional genome-alignment (e.g., LASTZ^16,17^) and mapping algorithms (e.g., LiftOver^18^). This apparent lack of enhancer conservation is largely due to rapid evolution of distal enhancers, but limitations in conventional computational genome alignment algorithms to detect conservation can also contribute significantly. Most conventional genome alignment algorithms utilize a *seed-and-extend* strategy, where short sequence matches are first obtained as *seeds* and then *extended* from both ends for further base pair alignment^16,17^. However, such nucleotide-level modeling of enhancer evolution may not be optimal for resolving sequence structures of enhancers, since enhancers are characterized by collections of multiple degenerate TFBS.

This paper addresses two major goals. First, we quantitatively derived 45 pairs of human and mouse cell/tissues from >1,000 ENCODE DNase-seq experiments^8,19–21^, through which we systematically quantified cell/tissue-specific enhancer conservation. Through multiple orthogonal analyses, we show that conservation levels of enhancers depend strongly on the cell/tissue type, which is partly explainable by association with transposable elements (TE). Second, we present a novel genome-alignment method that incorporates gapped-kmer features to model sequence degeneracy of enhancers: *gkm-align*. This feature choice is motivated by sequence modeling using gapped-kmer composition which have been shown^22^ to effectively represent biological sequences, accurately predict cell-specific enhancers, and discover regulatory vocabularies associated with TF binding (gkm-SVM)^23–25^. The effectiveness of this modeling agrees with the prevailing model that enhancers are defined by clusters of degenerate TFBS^23–26^. *Gkm-align* incorporates this idea and aligns human and mouse sequences by their gapped-kmer composition. Using enhancers of the 45 human/mouse cell/tissue pairs, we systematically evaluated the *gkm-align* algorithm, and discover thousands of novel conserved enhancers. Further, we show that the discovery rate of conserved enhancers can further be increased by incorporation of gkm-SVM derived cell-specific regulatory vocabularies, which we show are conserved between human and mouse.

## Results

### Cell-specific enhancer and promoter gkm-SVM regulatory vocabularies are conserved across mammals, while enhancers rapidly evolve

Enhancers are distal to transcription start sites (TSS) and harbor binding sites for transcription factors (TF). The cell-specific expression of these TFs leads to cell-specific enhancer chromatin accessibility and transcriptional regulation. Over the past decades, the ENCODE consortium has generated thousands of DNase-seq experiments^8,19–21^, across diverse human and mouse cell/tissues, and this comprehensive library of experiments has allowed us to robustly and systematically identify enhancer elements. Chromatin accessible peaks broadly fall into enhancer and promoter classes, which have different sequence features and conservation properties. To robustly define these classes, we use two genomic features: distance to the nearest TSS, and the cell-specificity of DNase I hypersensitivity (**Fig 1A**). We classified all DNase I hypersensitive sites (DHS) farther than 2 kilobases from the nearest TSS as *distal* (if not, *proximal*) and DHSs that are accessible in less than 30% of all biosamples in the ENCODE database (N_human_=1,270; N_mouse_= 153) as *cell-specific* (if not, *constitutive*). This partitions DHSs into four classes: distal cell-specific, proximal constitutive, distal constitutive, and proximal cell-specific DHSs (**Methods**).

**Fig. 1.**
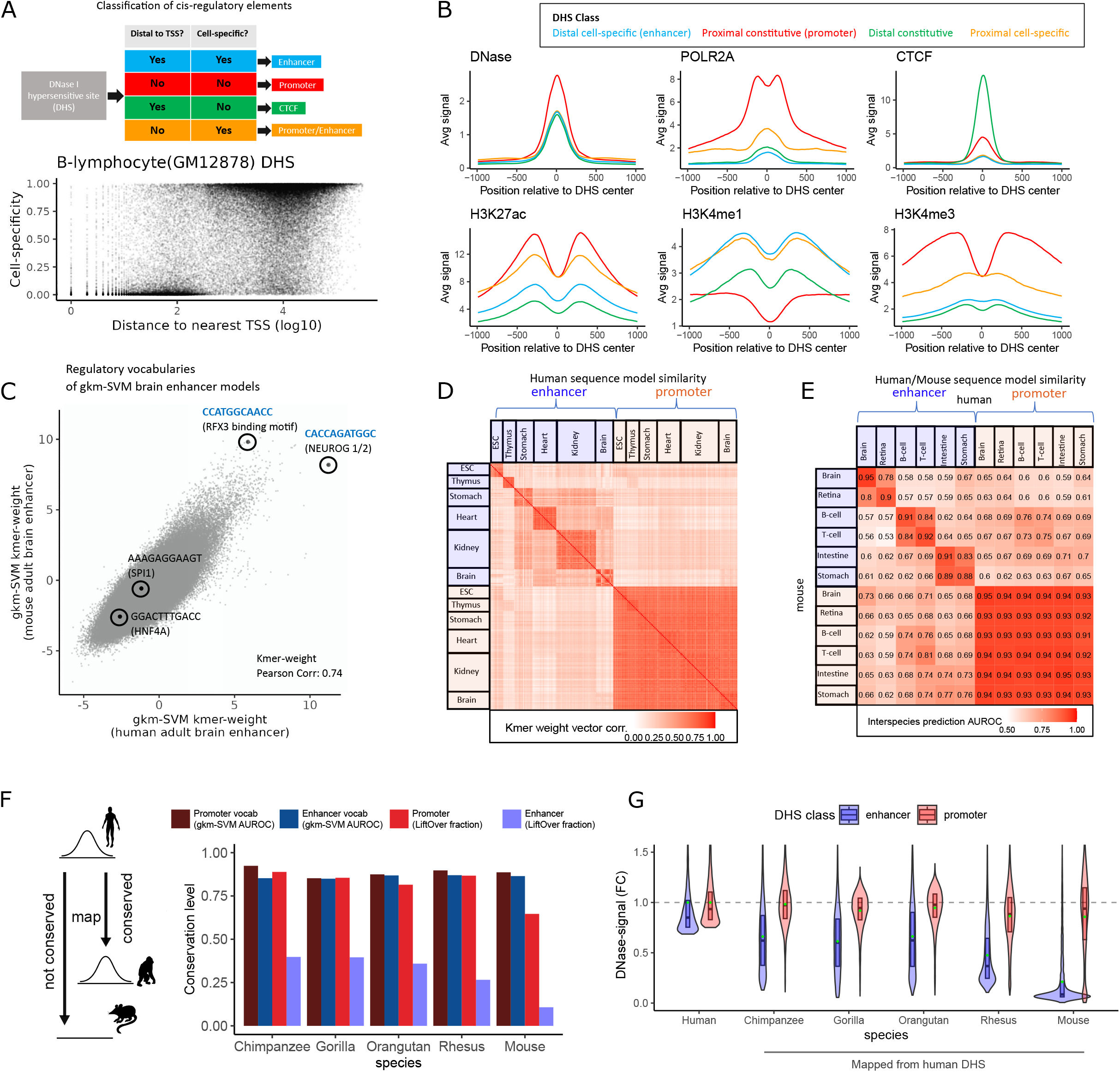
Cell-specific enhancer and promoter gkm-SVM regulatory vocabularies are conserved across mammals, while enhancers rapidly evolve. **A)** B-lymphocyte DHSs (N = 55,715) distance to nearest TSS and cell-specificity across 1,270 biosamples. **B)** Classification of B-lymphocyte DHSs by their distances to nearest TSS (proximal: < 2kb, distal: > 2kb) and cell-specificity (cell-specific: DHS in less than 30% of all biosamples, constitutive: otherwise); average epigenetic signals around DHS peak centers by DHS class. **C)** Pairwise comparisons of kmer-weight vectors, derived from enhancers (cell-specific distal DHS), for a pair of similar samples: human and mouse adult brain (N_kmer_ = 2,097,152; Pearson Corr. = 0.74). **D)** Pairwise comparisons of gkm-SVM kmer-weight vectors, by Pearson correlation, across various human embryonic biosamples (N_ESC_ = 11 N_Thymus_ = 9, N_Stomach_ = 15 N_Heart_ = 19, N_Kideny_ = 32, N_Brain_ = 14) trained on enhancers and promoters. **E)** Human-mouse interspecies DHS prediction accuracy (AUROC) across various cell-tissues (avg. between mouse element prediction using model trained on human elements and vice versa). **F)** Schematic of enhancer mapping and conservation level of human fibroblast enhancer/promoter regulatory vocabularies (gkm-SVM interspecies enhancer prediction AUROC) and enhancer/promoter CREs (fraction of conserved enhancers mappable by LiftOver) in chimpanzee, gorilla, orangutan, rhesus, and mouse. **G)** Distribution of DNase accessibility in human fibroblast enhancers and promoters (top 10,000 in each DHS class by total DNase-seq reads mapped) and distribution of DNase-signal at primate and mouse loci mapped from human enhancers and promoters. Signals are normalized as fold changes from average fibroblast enhancers and promoter accessibilities in respective species (top 10,000 in each DHS class by DNase-seq read mapped).

We observed distinct biochemical signatures in these four element classes. Distal cell-specific DHSs show strong markers of enhancer activity such as ChIP-seq signal for H3K4me1 and lack markers of promoter activity (POLR2A, H3K4me3) (**Fig 1B, Supp Fig 1-2**) (**Methods**; ENCODE experiment IDs listed in **Supplementary Table 1**). By contrast, proximal constitutive DHSs have the highest level of chromatin accessibility among the four classes and displayed clear signatures of promoter activity (POLR2A, H3K4me3) and depletion of enhancer marks (H3K4me1). These classification criteria allowed us to robustly define enhancer elements without the need for diverse histone ChIP-seq experiments, which are currently unavailable for many biosamples assayed with the DNase-seq experiments. Further, many distal constitutive DHSs appear to be CTCF binding sites (a known regulator of chromatin topological organization), and proximal cell-specific DHSs show mixed signatures of enhancers and promoters, which emphasizes the utility of the two criteria (TSS distance; cell-specificity of chromatin accessibility) for precisely defining enhancer elements (**Fig 1A-B**). For the rest of the paper, we will refer to distal cell-specific DHSs as enhancers and proximal constitutive DHSs as promoters.

**Fig. 2.**
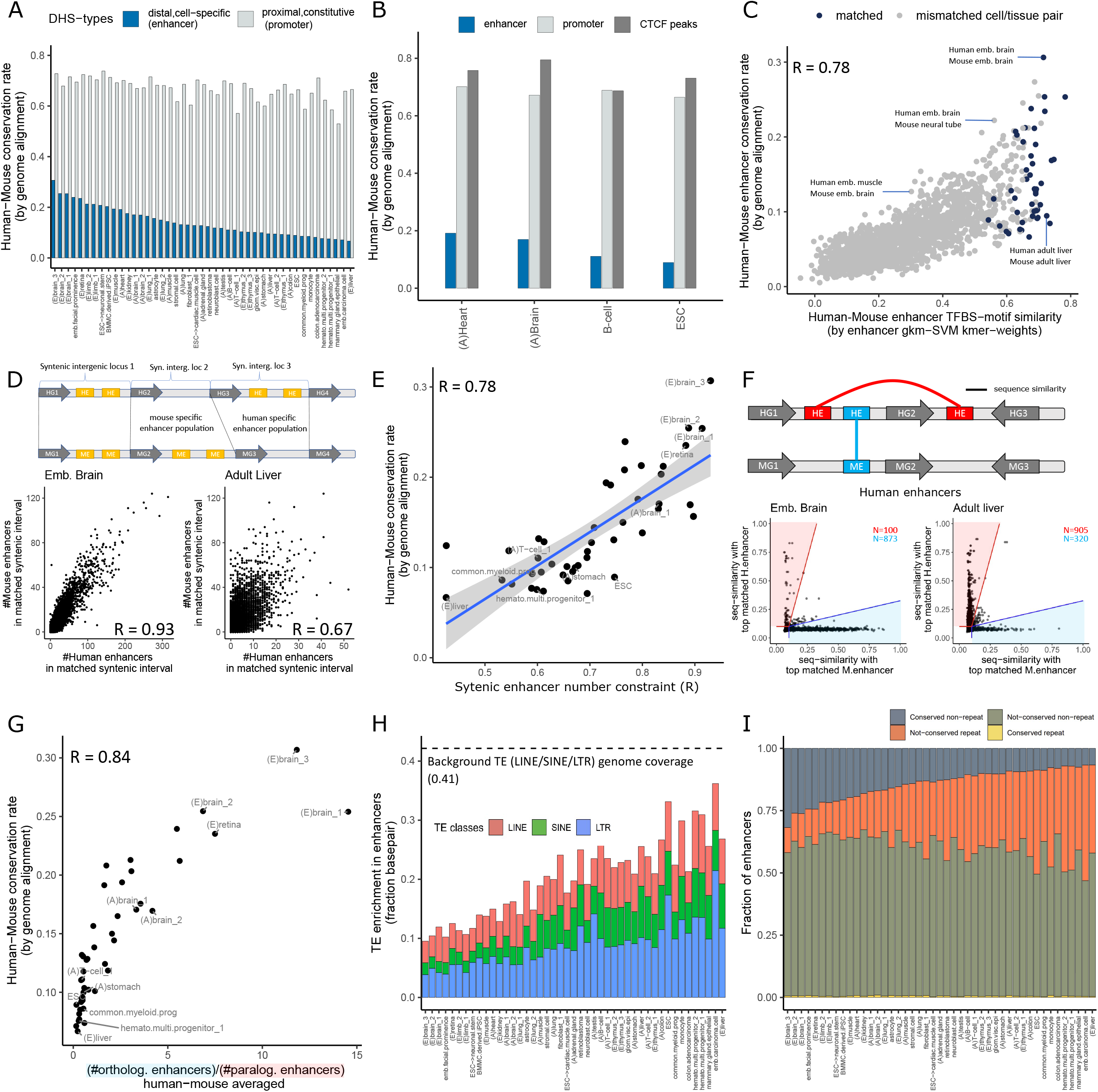
Enhancer conservation levels are highly variable across cell types. **A)** Human-mouse enhancer (distal cell-specific DHS) and promoter (proximal constitutive DHS) conservation rates (mappability to DHS by LASTZ/LiftOver genome alignment/mapping algorithms in orthologous syntenic intergenic loci. Mean of human to mouse and mouse to human mappings) across the 45 human-mouse cell-tissue pairs (A: adult, E: embryonic); sorted by enhancer mappability (identical orders for **Fig 2H-I**). **B)** Human-mouse conservation rate for enhancers, promoters, and CTCF ChIP-seq peaks for B-cell, brain, ESC, and heart. **C)** Human-mouse enhancer conservation rate by alignment (mappability by LASTZ/LiftOver) vs. human-mouse enhancer regulatory vocabulary conservation rate (correlation of gkm-SVM kmer-weights) for pairs of orthologous gkm-SVM matched tissues (e.g., human and mouse brain; N = 45) and mismatched cell/tissues (e.g. human T-cell and mouse brain; N = 1,980) (R =.78) **D)** Schematics of human-mouse syntenic intergenic loci (HE: human enhancer; ME: mouse enhancer; HG: human gene; MG: mouse gene); Comparison of human and mouse enhancer numbers in matched syntenic intergenic loci (N = 12,704) for embryonic brain (syntenic enhancer number constraint R = .91) and adult liver (R = .67) **E)** Syntenic enhancer number constraint vs. human-mouse enhancer conservation rate (N = 45, R = .78; linear regression line and 95% confidence interval). **F)** Schematic describing how orthologous and paralogous enhancers are defined (HE: human enhancer, ME: mouse enhancer, G: gene; line indicates sequence homology); dots represent 5,000 human enhancers w/ highest DNase signal. Gapped-kmer sequence similarity with top matched mouse enhancers vs with top matched human enhancers. Enhancers in red shaded region are classified as paralogous enhancers; enhancers in blue shaded regions are classified as orthologous enhancers. **G)** Ratio of orthologous to paralogous enhancers across the 45 cell/tissue pairs (avg. of human and mouse) vs. human-mouse enhancer conservation rate (by genome alignment) **H)** Total fraction of enhancer base pairs annotated by each class of transposable elements (avg. of human and mouse). **I)** Fraction of conserved enhancers and overlap with TEs (repeat = enhancers with more than 50% base pair overlap with type I TEs; non-repeat = enhancers with zero overlap with any repeat annotations).

Enhancer regulatory vocabularies, obtained through gkm-SVM training on enhancers, are cell-specific. Gkm-SVM is a sequence-based machine learning method that learns to effectively distinguish enhancers from inactive genomic elements by learning the weighted combination of gapped-kmers that predict enhancers^23^. This information can be summarized into a kmer-weight vector, where kmers comprised of predictive gapped-kmers – or enhancer vocabularies– are assigned higher weights. For easier interpretation, we can map gapped-kmers to kmer weights^23^. The biological relevance of enhancer regulatory vocabularies has been demonstrated by their utility in predicting functional impacts of enhancer sequence variants^24–28^. All human and mouse gkm-SVM models used in this study are publicly available in the ENCODE portal, and their aliases and experimental inputs are listed in **Supplementary Table 2-3**.

Enhancer kmer-weight vectors encode DNA binding motifs of core TF regulators that determine cell/tissue identity, leading to similarities of enhancer regulatory vocabularies by cell/tissue types. For example, gkm-SVM models trained on human and mouse adult brain enhancers detect the same TFBS, encoded by highly similar enhancer regulatory vocabularies (**Fig 1C**; R = 0.74). For example, one of the top predictive kmers (CACCAGATGGC) shared between human and mouse brains is a TFBS for neurogenic TFs NEUROG1/2 and ATOH1^29–31^. Another predictive kmer for both human and mouse (CCATGGCAACC) is bound by RFX family TFs; of which RFX3/RFX5/RFX7 are highly expressed in the human brain^32^, and their mutations are linked to intellectual and behavioral abnormalities^33^. On the other hand, kmers associated with TFs highly expressed in non-neural cell/tissues, such as AAAGAGGAAGT (SPI family; blood cell development^34^) and GGACTTTGACC (HNF4A; liver/pancreas^35^), have low weights in both human and mouse brain enhancer models. Cell/tissues of distinct identity have low similarity in enhancer kmer weight vector (**Supp Fig 3**; R_brain vs monocyte_, R_spinal cord vs macrophage_, R_brain vs liver_ = 0.085, 0.11, 0.19). Across a wider range of cell/tissues, biosamples of the same cell/tissue identity consistently show high kmer-weight correlation (mean R = 0.72) while pairs of kmer-weights from distinct cell/tissues show lower correlation (mean R = 0.31) (**Fig 1D**). By contrast, kmer-weights obtained from promoters show low cell-specificity, having high kmer-weight correlation for all pairs of cell/tissues (same tissue mean: 0.80; distinct tissue mean: 0.75). This is consistent with past observations that enhancers are bound by TFs with cell-specific expression while promoter binding TFs are relatively less cell-specific^36^.

**Fig 3.**
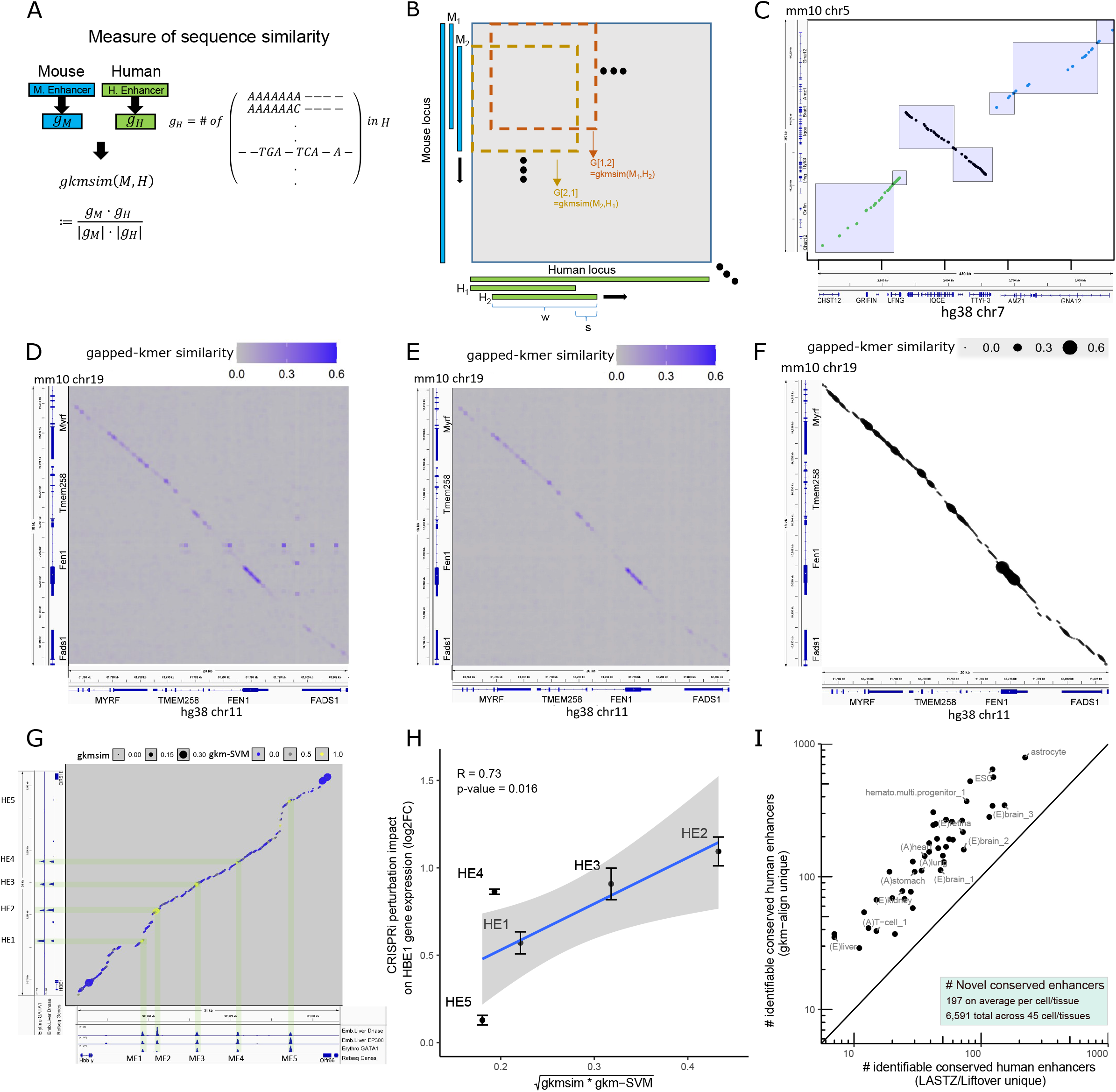
gkm-align algorithm identifies conserved enhancers by finding optimal alignment path of maximum gapped-kmer similarity. **A)** Sequence similarity between a mouse (M) enhancer and a human (H) enhancer is quantified by their similarity in gapped-kmer compositions (gapped-kmer similarity, or gkm-sim). **B)** Schematic describing computation of pairwise gkm-similarity of all pairs of sliding windows in syntenic genomic loci of the two species. The pairwise similarity values are encoded in a gapped-kmer similarity matrix *G.* **C)** Schematic describing how whole-genome alignment is performed using the GNA12 inversion locus as an example (dots: short sequence matches. colors: groups of short matches in syntenic blocks; boxes: pairs of human/mouse syntenic loci from which gkm-similarity matrices derive). Visualization of gapped-kmer similarity matrix (*G*) in FADS syntenic locus **D)** without gkm-SVM repeat masking and **E)** with masking. **F)** Identification of colinear series of conserved elements using matrix *G*. **G)** Alignment of the HBB Locus Control Region (dot size: gkm-similarity; color: gkm-SVM prediction score at corresponding human locus using gkm-SVM model trained on mouse embryonic liver enhancers. Highlights: CREs); HE: human element; ME: mouse element. **H)** Combining gkm-similarity of (HE, ME) and gkm-SVM score (of HE using mouse embryonic liver enhancer trained model) to predict regulatory activities of human elements (HE) measured by CRISPRi perturbation (R=0.73; error bars: minimum and maximum of 2 bio replicate effect sizes) (linear regression line and 95% confidence interval; p-value (t): 0.016) **I)** Number of human-mouse conserved enhancers that are identifiable uniquely by LASTZ/LiftOver (x-axis) and gkm-align (y-axis) for each of the 45 cell/tissue pairs. Gkm-align identifies conserved enhancers missed by LASTZ/LiftOver in all tissues. Relative to LASTZ/LiftOver, gkm-align discovers 197 novel enhancers on average per cell/tissue and total 6,591 novel enhancers across all 45 cell/tissues.

Comparing all human and mouse tissue pairs, we find that enhancer regulatory vocabulary is conserved, as shown for a subset of cell/tissues in **Fig 1E**. We trained gkm-SVM enhancer prediction models for a set of human and mouse biosamples (brain, retina, B-cell, T-cell, intestine, stomach), and evaluated whether gkm-SVM regulatory vocabularies trained on one species are predictive of enhancers in the other species (i.e., train on human enhancers, predict on mouse enhancers & train on mouse, predict on human) (**Methods**). We evaluated the similarity of human and mouse models by taking average between reciprocal interspecies prediction accuracy (AUROC). For matched cell/tissues, accuracy of interspecies enhancer prediction is high (N = 6; mean AUROC = 0.91), while prediction accuracy between distinct cell/tissues is low (N = 30; mean AUROC = 0.65). On the other hand, interspecies prediction accuracies for promoters is high for both mismatched (e.g., human brain & mouse B-cell; N = 30; mean AUROC = 0.93) and matched cell/tissues (N = 6; mean AUROC= 0.94). These results are consistent with past observations that expression of core TFs^37,38^ and their DNA binding affinities^15^ are well-conserved between human and mouse.

Although enhancer vocabularies are highly conserved, mapping orthologous enhancers conserved between distant mammals remains computationally challenging. To assess enhancer conservation rate, we will use as a metric the fraction of predicted enhancers in one species that is also chromatin accessible in the other species (**Fig 1F**). To demonstrate the challenge of mapping enhancers, we will first use a conventional mapping method (LASTZ/LiftOver) and compute the conservation rate of human fibroblast enhancers and promoters using a set of DHS data in fibroblasts from diverse mammals^39^ (chimpanzee, gorilla, orangutan, rhesus, and mouse^8^, in increasing divergence from human) (**Methods**; **Fig 1F-G**). About 40% of enhancers are mappable between human and chimpanzee, and this value rapidly decreases to 11% in human and mouse as evolutionary distance to human increases (72% decrease in conservation rate; **Fig 1F**). Promoter conservation rate also decreases from 89% between human and chimpanzee to 65% between human and mouse but at slower rate of 27%. In contrast, regulatory vocabularies of both enhancers and promoters as quantified by gkm-SVM are constant across all species (**Fig 1F**); interspecies prediction accuracy (AUROC) of enhancers and promoters between human and mouse (AUROC enhancer: 0.86; promoter: 0.89) are as high as the prediction accuracy between human and chimpanzee (AUROC enhancer: 0.85; promoter: 0.92). As an alternative metric, we can count the read signal in the orthologous loci relative to average elements in that class. This confirms the rapid reduction in enhancer conservation rate (**Fig 1G**), where we show, for each species, distributions of DNase-signal at orthologous loci mapped from human fibroblast enhancers and promoters (**Methods**). Orthologous chimpanzee loci mapped from human enhancers and promoters respectively are on average 66% and 98% as accessible as average chimpanzee enhancers and promoters. These signals decrease dramatically to 21% (enhancer) and 86% (promoter) when we map human enhancers and promoters to mouse, further underscoring the rapid evolution of enhancers.

We can use enhancer regulatory vocabulary to identify the most similar pairs of human and mouse cell/tissues for further quantitative assessment of conservation (**Fig 2**). We generated a comprehensive list of 45 human/mouse cell/tissue pairs matched by gkm-SVM enhancer vocabularies (**Supplementary Table 4**; **Methods**). We will also use this set of 45 pairs of cell/tissues to evaluate our novel genome-alignment method, *gkm-align*, against conventional methods (**Fig 3-4**).

**Fig 4.**
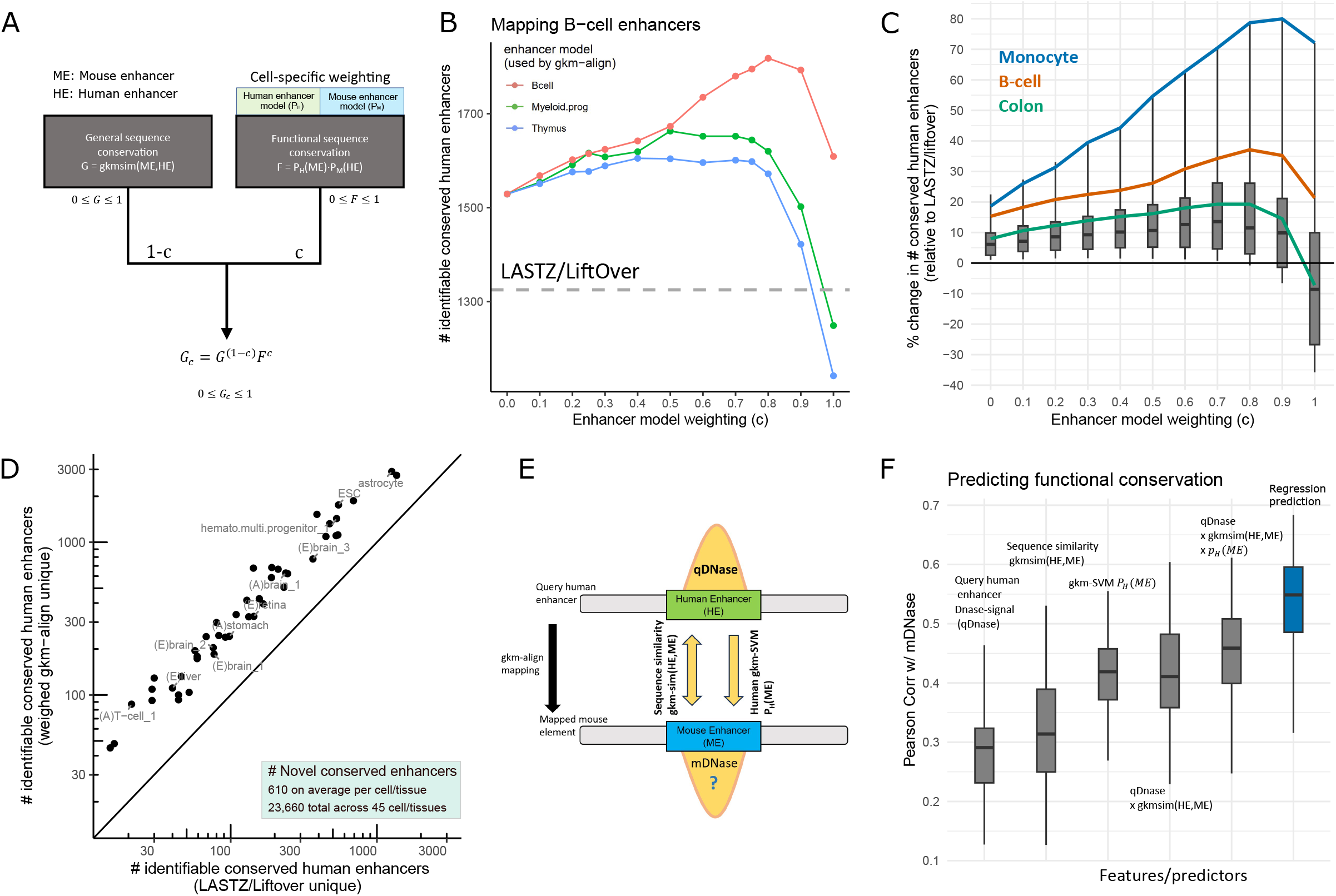
gkm-align identifies more novel conserved enhancers and robustly predicts functional conservation when combined with cell-specific information. **A)** Schematic describing how cell-specific gkm-SVM enhancer prediction model is incorporated into *gkm-align* for cell-specific weighted alignment. **B)** Number of B-cell human enhancers mappable to mouse B-cell enhancers using LASTZ/LiftOver (grey dashed line) and *gkm-align* weighted by gkm-SVM enhancer models trained on B-cell (red), myeloid progenitor cell (green), and thymus enhancers (blue) with varying weights **C)** percent change in the number of identifiable human/mouse enhancer pairs by gkm-align, relative to LASTZ/LiftOver, across 45 cell-tissue pairs using cell-type matched gkm-SVM model weighting. Line plots for a subset of cell/tissues. **D)** Number of human-mouse conserved enhancers that are identifiable uniquely by LASTZ/LiftOver (x-axis) and gkm-SVM weighted cell-specific gkm-align (y-axis) for each of the 45 cell/tissue pairs. Relative to LASTZ/LiftOver, weighted gkm-align discovers 610 novel enhancers on average per cell/tissue and total 23,600 novel enhancers across all 45 cell/tissues. **E)** Schematic describing regression model for ranking enhancer mapping to mouse (mDNase) in terms of human enhancer features (qDNase = DNase-signal at query human enhancer; gkm-sim = gapped-kmer sequence similarity between human and mouse element; P_H_(M) = human gkm-SVM score of mapped mouse element). **F)** Predicting DNase-seq signal at mouse loci mapped from human enhancers (mDNase) using combinations of features described in Fig. 4E across the 45 human/mouse cell/tissue pairs.

### Enhancer conservation levels are highly variable across cell types

In the 45 pairs of human and mouse cell/tissues, we observed intriguingly high cell/tissue-specific variability in enhancer conservation. As in **Fig 1G**, we defined the conservation rate of human enhancers as the fraction of human enhancers that map to mouse DHS of the matched cell/tissue (e.g., human brain/mouse brain), constraining the mapping by LASTZ/LiftOver to syntenic intergenic loci (**Methods**). Conservation rates of mouse enhancers were defined similarly, and human-mouse conservation levels were defined as the average of the two reciprocal directions. Promoters showed consistently high conservation rate across the 45 tissues (mean 67%) while enhancers showed highly variable conservation rate, ranging from 6.7% (embryonic liver) to 31% (embryonic brain) (**Fig 2A**). Such lack of conservation is not observed in CTCF ChIP-seq peaks, which are also often distal to TSSs (conservation rate for brain, heart, B-cell, ESC = 82%, 79%, 72%, 76%; **Fig 2B**), suggesting that many CTCF loops and topologically associated domains are conserved^14,40^. The strong and somewhat counter-intuitive tissue specificity of enhancer conservation will be explored extensively below.

We observed low enhancer conservation rates in some tissues in spite of the fact that their regulatory vocabulary is highly similar between human and mouse. To explore this systematically, we computed the similarity of enhancer regulatory vocabularies (measured as Pearson Corr. of enhancer kmer-weight vectors), for both matched and mismatched pairs of human and mouse cell/tissue pairs (e.g., matched: human brain & mouse brain; mismatched: human brain & mouse muscle; N_matched_ =45; N_mismatched_ = 45^2^-45=1,980; **Fig 2C**). Overall, conservation rates of enhancers and similarity of enhancer regulatory vocabularies correlate highly (R = 0.78, **Fig 2C**), indicating that human and mouse cell/tissue that share similar sets of core TF regulators also tend to share a higher number of orthologous enhancers. While enhancer regulatory vocabularies were overall highly similar for all matched human/mouse cell/tissue pairs (relative to the mismatched pairs), interestingly, their enhancer conservation rates varied widely (vertical spread of black points). For example, the human/mouse adult liver pair had almost identical similarity of enhancer regulatory vocabulary (R = 0.72) as human/mouse embryonic brains (R = 0.71), but the adult liver enhancer conservation rate was 9.4% (less than 1/3 of the embryonic brain), even lower than the enhancer conservation rate between a mismatched pair of human embryonic muscle and mouse embryonic brain (12.7%). This implies that some cell/tissues, while maintaining their core TF regulators and their DNA binding specificities, have experienced more incidence of enhancer turnover than other cell/tissues.

To eliminate the possibility that the highly cell/tissue specific rate of enhancer conservation is a bias of the LASTZ/LiftOver alignment/mapping algorithms, we performed an orthogonal analysis. Using 12,455 syntenic intergenic loci of human and mouse derived from 15,500 orthologous protein coding genes^20^ (**Supplementary Table 5**), we simply counted the number of human and mouse enhancers located in each of the matched syntenic intergenic loci (**Fig 2D; Methods**). We compare the correlation between the number of human and mouse enhancers in respective syntenic intergenic loci, which we will refer to as “syntenic enhancer number constraint,” and it imposes an upper limit for mappability of human/mouse enhancers in the matched syntenic loci. If the number of enhancers in syntenic intervals is not conserved, there is no way the enhancers can be conserved at the sequence level unless they arose through duplication. Embryonic human/mouse brain, which had the highest rate of enhancer conservation, also showed the highest level of syntenic enhancer number constraint (R=0.93; **Fig 2D**), while adult human/mouse liver had lower syntenic enhancer number constraint (R=0.67), with occasionally drastically different numbers of enhancers in matched syntenic intergenic loci. This is consistent with reports of species-specific rewiring of transcription in the liver^41,42^ and a relatively slower rate of transcriptomic divergence of the brain across mammals^43^. The lack of syntenic enhancer number constraint appears in cell/tissues with low enhancer conservation level (**Fig 2E**), and hints that the lower rate of conserved enhancers in some cell/tissues, as predicted by genome alignment, is an inherent property of the regulatory landscape.

This apparent lack of syntenic enhancer number constraint is largely driven by species-specific enhancer duplication, where cell/tissues with lower level of enhancer conservation tend to have a higher proportion of paralogous enhancers. To estimate the proportion of orthologous and paralogous enhancers in each human cell/tissue (in the context of human-mouse common ancestry), we labeled a human enhancer as paralogous if it has high sequence homology with another human enhancer but lacks sequence homology with any mouse enhancer, and similarly labeled it as orthologous if it has high sequence homology with a mouse enhancer but lacks homology with any other human enhancers (**Fig 2F**; **Methods**). Based on this criterion, of the 5,000 human embryonic brain enhancers with highest DNase I accessibility, 873 were identified as orthologous and 100 as paralogous enhancers (ratio: 8.73). In contrast, adult liver had 320 and 905 orthologous and paralogous enhancers (ratio: 0.35). These estimated ratios of orthologous to paralogous enhancers, averaged between human and mouse, closely matched with the syntenic enhancer number constraint and with enhancer conservation rate for the 45 cell/tissue pairs (**Fig 2G**; **Supp Fig 4**), indicating that enhancer duplication events have been a significant contributor to the divergence in enhancer landscapes.

These duplication events are largely driven by transposable elements (TE). The paralogous enhancers show significant enrichment of LTR retrotransposons across diverse cell/tissues (**Supp Fig 5**; **Methods**), and we observed a general trend that the cell/tissue pairs with low enhancer conservation level tend to have high enrichment of transposable elements (**Fig 2H** – bars in the same order as **Fig2A**; **Supp Fig 9A**; R = -0.88). Quantifying TE enrichment as the fraction of total enhancer base pairs that overlap with a TE, we observed that embryonic brain enhancers, averaged between human and mouse, had less than 10% enrichment of type I transposable elements (LTR/LINE/SINE) while some cell/tissues, such as ESC, had TE enrichment as high as 33%. Overall, TE enrichment in enhancers appears lower than its genome-wide coverage (>40%), with SINE elements especially depleted in enhancers of most cell/tissues (**Supp Fig 6 E-F**). However, interestingly LTR elements appear to be highly enriched in enhancers across multiple cell/tissues, and their enrichment grows in enhancers with the strongest DNase I accessibility (**Supp Fig 6 A-B**). Although we do observe clear signals of DNase I accessibility in LINE elements (at its 5’ end) for multiple cell/tissues (**Supp Fig 7**), these generate weak DNase I peaks, and LINE elements are depleted in enhancers of most cell/tissues (**Supp Fig 6 C-D**). Like LTR, LINE enrichment increases with increasing DNase I accessibility, surpassing the genomic average for subsets of top 1,000 enhancers of human colon and ESC with the highest level of DNase I accessibility (**Supp Fig 6 C-D**). By contrast, SINE elements are more depleted in enhancers with higher DNase-I accessibility. This TE-specific and cell/tissue-specific variation in TE-enhancer association suggests that TEs may have a functional role in shaping the enhancer landscape^44–48^, but it is difficult to separate function from their naturally increased tendency to transpose into accessible regions.

**Fig 5.**
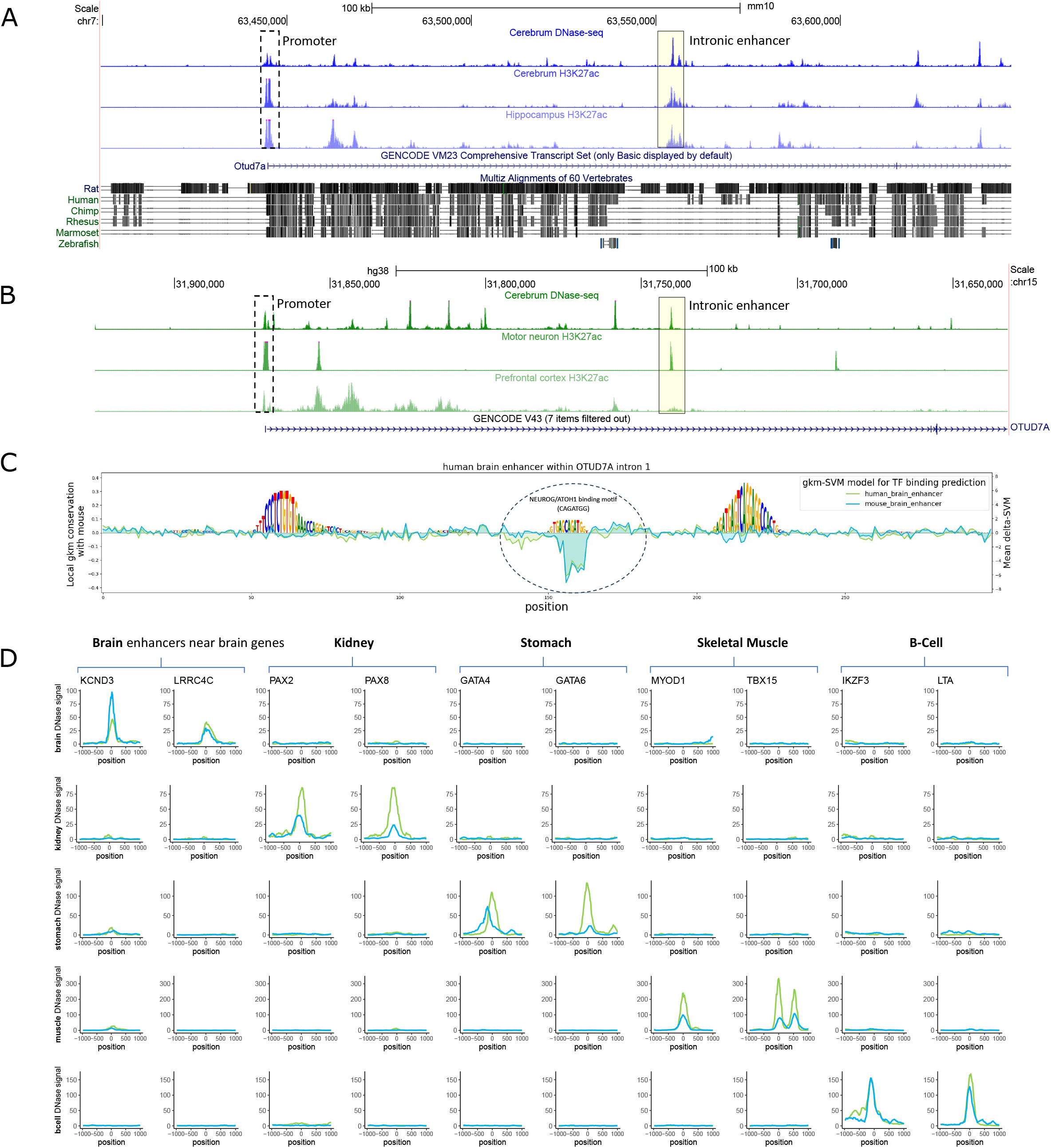
Examples of novel enhancers from expanded catalogue of human/mouse orthologous enhancers. **A)** Genome browser visualization of mouse and **B)** human OTUD7A loci. Yellow boxes indicate the identified conserved enhancers. **C)** Visualization of conserved binding sites in human OTUD7A intronic enhancer by sequence conservation with the orthologous mouse enhancer. Logo height represents local gapped-kmer sequence conservation score with mouse, and line plots indicate TF-binding prediction by delta-SVM models trained on human or mouse brains enhancers. **D)** Visualization of DNase accessibility of orthologous enhancers from the five distinct cell/tissues (Green: human, Blue: mouse; position relative to DHS center of human/mouse orthologous enhancers; signal: fold change from genomic average).

Most enhancers with overlapping TE annotations are species-specific (**Fig 2I**; **Methods**), and the increase in TE enrichment in enhancers explains much of the decrease in enhancer conservation across cell/tissues. This is not limited to weaker DHS, and in fact TE enrichment in enhancers grows with DNase-I accessibility. We show this by repeating the analysis of **Fig 2I** using only the top 1,000 enhancers with the highest DNase-signal (**Supp Fig 10**). In summary, enrichment of TE-associated enhancers contributes heavily to the observed cell/tissue dependent variability in enhancer conservation (orange bars in **Fig 2I** and **Supp Fig 10** grow with decreasing conservation). However, intriguingly, we also observe a decreasing trend of conservation in non-TE associated enhancers as TE-enrichment in cell/tissues increases (grey bars in **Fig 2I** and **Supp Fig 10** shrink with decreasing conservation; **Supp Fig 9B**; R = -0.84). We find it curious that the increased enhancer duplication driven by TEs is correlated with the reduction in the conservation level of non-TE associated enhancers. It is possible that TE driven functional redundancy allows more rapid evolutionary turnover in these tissues.

### *gkm-align* algorithm identifies conserved enhancers by finding the alignment path of maximum gapped-kmer similarity

Identification of conserved enhancers in evolutionarily distant mammals, such as mice, is made difficult by their rapid evolution; and limitations of genome alignment algorithms may underestimate conservation. We do not know how significantly LASTZ/LiftOver alignment inaccuracy contributes to the low rates of conservation shown in **Fig 2**. To address this issue, we next present a novel genome alignment algorithm that differs from previous methods by using gapped-kmer sequence features that more readily capture the functional elements of enhancer sequences.

Enhancers contain degenerate clusters of TFBS, and enhancer mutagenesis studies have shown that enhancer function is strongly affected by mutations within binding sites and robust to mutations between binding sites^28,49^. This modular architecture more readily tolerates insertions/deletions between TFBS and small structural variations. To exploit this modular structure, *gkm-align* uses a sequence similarity metric that compares a pair of sequences (e.g., width of 300 base pairs) by their gapped-kmer composition (**Fig 3A; Supplementary information** for algorithmic details). Gapped-kmers are generalizations of kmers; they contain a fixed number of gaps, which represent any nucleotide, and model degenerate positions in TFBS. There exist 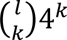 gapped-kmers with size *l* and *k* non-gapped positions (e.g., N=5,406,720 for *l*, *k*=11, 7), and the gapped-kmer similarity (*gkm-sim*) for a pair of sequences is computed as cosine similarity of vectors, each encoding the counts of gapped-kmers in the respective sequence.

To align a pair of human/mouse loci, we first compute a gkm-similarity matrix (*G*) of all pairs of sliding window subsequences of the human and mouse loci (**Fig 3B**; **Supp info B**). The size of this matrix will depend on the locus size; for example, human/mouse loci of 20 kilobases (e.g., FADS loci) have ∼1,000 subsequences of 300 base pair windows sliding by 20 base pairs, and the gkm-similarity matrix (*G*) of dimension 1,000 x 1,000 encodes all pairwise gkm-similarities of the human and mouse subsequences (as shown in **Fig 3D**). Exons of matched orthologous genes in human and mouse show highest levels of sequence similarity, but regions of low complexity (e.g., tandem repeats) also show high interspecies sequence similarity due to their prevalence and uniform sequence composition (identifiable as a row of horizontal dots in **Fig 3D**). To remove these repetitive sequence matches, we train gkm-SVM to detect and mask sequence patterns that are ubiquitous across the human and mouse genomes (**Fig 3E; Supp info E.3**). We then compute *G* using the masked sequences, to which we apply a variant of Smith-Waterman algorithm to identify an optimal alignment path that encodes how human/mouse loci diverged (**Fig 3F; Supp info C**). This method of alignment is extended genome-wide by utilizing orthologous gene annotations and short sequence matches (**Fig 3C; Supp info D**).

We next demonstrate the *gkm-align* algorithm at the well-studied hemoglobin beta (HBB) locus control region (LCR) ^50^. These loci in human and mouse each contains 4-5 enhancers, and the human enhancers have shown to be capable of regulating mouse HBB expression through transgenic mouse experiments^51^. We aligned the HBB LCRs of human and mouse using gkm-align and mapped the five mouse enhancers to human (**Fig 3G**). The five mouse enhancers (labeled as ME1, …, ME5) all have strong DNase I accessibility and EP300 binding in mouse embryonic liver (where HBB is active) and are bound by GATA1^52^ in erythroblasts; however only four of the five human loci mapped from these mouse enhancers (labeled as HE1, …, HE5) show strong marks of enhancers. HE5 has weak GATA1 binding in erythroblast and low DNase I accessibility in embryonic liver. Further, inhibiting HE5 with CRISPRi has the weakest effect on downregulating HBE1 expression among the five putative human enhancers mapped from mouse^53,54^ (**Fig 3H**; **Methods**). It is likely that HE5 may have accumulated mutations leading to loss of regulatory activity. This loss of regulatory activity is also predictable by gapped-kmer sequence similarity metrics. As will be justified further below, **Fig 3H** shows that the geometric mean of gapped-kmer sequence similarity and interspecies gkm-SVM enhancer prediction is consistent with the CRISPRi effect at HBB enhancers (R=0.73).

We applied *gkm-align* genome-wide across the 45 cell/tissue pairs and identified many novel conserved human enhancers which are not predicted by LASTZ/LiftOver (either predicted to be deleted or which LiftOver maps to inactive regions). For all the 45 cell/tissue pairs, gkm-align mapped a higher number of human enhancers to mouse enhancers than LASTZ/LiftOver, with the increase in enhancer mappability ranging from 1% (embryonic limb) to 22% (hematopoietic multipotent progenitor cells) (**Fig 3I**, **Fig 4C**). For the cell/tissue pair of human hippocampus astrocyte and mouse Müller cells (both of which are glial cells), *gkm-align* successfully mapped 791 human enhancers to mouse enhancers (cell-specific distal DHS peaks, **Methods**), which are incorrectly mapped by LASTZ/LiftOver (either deleted or map to inactive mouse regions). Conversely, only 222 human enhancers were correctly mappable uniquely by LASTZ/LiftOver but incorrectly mappable by *gkm-align*. 8,559 human glial enhancers were mapped to mouse enhancers by both methods. Together across the 45 cell/tissues, gkm-align identified 6,591 novel conserved enhancers. This greatly increases the number of human enhancers which can be functionally tested for disease relevance in mouse models.

### *gkm-align* identifies additional novel conserved enhancers and robustly predicts functional conservation when combined with cell-specific information

Although *gkm-align* outperforms LASTZ/LiftOver, the sequence similarity metric does not explicitly make use of cell/tissue specific regulatory vocabulary. Gkm-SVM enhancer regulatory vocabularies, encoding TFBS motifs, are well conserved between human and mouse (**Fig 1**), and they can be incorporated into gkm-align both to improve discovery of conserved enhancers and to quantify their functional conservation. This additional information leads to an expanded catalog of human enhancers testable through mouse models, ranked by likelihood of conserved regulatory roles.

We incorporate cell-specific gkm-SVM regulatory vocabularies into gkm-align following a simple and intuitive model: if a pair of human and mouse enhancers, denoted as HE and ME, are orthologous, then they should have similar DNA compositions (i.e., general sequence conservation), and should both contain conserved TFBS motifs relevant to the shared cellular context (i.e., functional sequence conservation) (**Fig 4A**). General sequence conservation (*G*) is quantified using gkm-similarity, as previously described. Functional sequence conservation (*F*) is computed using interspecies gkm-SVM prediction scores, which we normalize to vary between 0 and 1 for interpretability (**Supp info E**). Denoting *P*_*H*_(*ME*) as normalized prediction score of a mouse element by a gkm-SVM model trained on human enhancers, we can interpret *P*_*H*_(*ME*) as the probability that *ME* can function as an enhancer in the orthologous human cellular context. *P*_*M*_(*HE*) is defined similarly (M: mouse; HE: human enhancer). Then, functional sequence conservation, computed as *F* = *P*_*H*_(*ME*) · *P*_*M*_(*HE*), can be interpreted as the probability that *ME* and *HE* can both function as enhancers interchangeably in human and mouse cellular contexts. For cell-specific weighted alignment, we combine the two measures of enhancer conservation into *G*_*c*_ = *G*^1−*c*^ · *F*^*c*^ (0 ≤ *c* ≤ 1), which adjusts the alignment path toward human/mouse element pairs with both similar sequence composition (*G*) and functional similarity in common cellular context (*F*).

Cell-specific weighted alignment by *gkm-align* identifies the highest number of conserved enhancers when it is combined with gkm-SVM enhancer prediction model trained on enhancers of relevant cell/tissue type. For example, LASTZ/LiftOver and gkm-align each identify 1,325 and 1,529 conserved human B-cell enhancers, but if gkm-align is combined with B-cell trained gkm-SVM enhancer prediction models, the number of identifiable conserved enhancers increases up to 1,818 at cell-specific enhancer model weighting parameter (*c*=0.8) (a 37% increase from LASTZ/LiftOver) (**Fig 4B**). The identification rate also increases when *gkm-align* is combined with gkm-SVM models of similar cell-types (with overlapping TFs), such as myeloid progenitor cells and thymus, each with peaks at 1,663 (*c*=0.5) and 1,605 (*c*=0.4) conserved enhancers. Similarly for colon enhancers, LASTZ/LiftOver, unweighted gkm-align (*c*=0), and cell-specific gkm-align (*c*=0.75) each identifies 1,125, 1,221, and 1,352 conserved enhancers, which corresponds to a 7.9% and 20% increase over LASTZ/LiftOver for unweighted (*c*=0) and weighted (*c*=0.75) gkm-align respectively (**Supp Fig 11**). Cell-specific weighting improves the identification rate of conserved enhancers for all pairs of 45 cell/tissues (**Fig 4C**). At *c*=0.9, we observe up to an 80% increase in conserved enhancer discovery over LASTZ/LiftOver for monocytes, with 16 cell/tissues with greater than a 20% increase for *c*=0.7 and c=0.8. A subset of cell/tissues exhibited limited improvement through gkm-SVM weighting (e.g., brain), but their identification rates remained higher than both unweighted gkm-align and LASTZ/LiftOver at *c*=0.5. Across the 45 cell/tissues, weighted cell-specific gkm-align discovers several hundred novel enhancers in every tissue and 23,660 total novel conserved enhancers across all 45 cell/tissues (**Fig 4D**). It should also be noted that, although cell-specific alignment discovers a higher number of conserved enhancers compared to unweighted *gkm-align*, there are also handful of enhancers identifiable by unweighted gkm-alignment but missed by cell-specific gkm-alignment (**Supp Fig 12**). These tend to be conserved enhancers with high general sequence similarities but with more degenerate TFBS. For this reason, both generic and cell-specific *gkm-align* should be used in parallel to maximize the span of identifiable conserved enhancers, and we provide genome-wide enhancer mappings for both methods.

Cell-specific information from gkm-SVM enhancer prediction models can also be combined with the gapped-kmer based sequence similarity metric (*gkm-sim*) to quantify functional conservation of orthologous enhancer pairs discovered through *gkm-align*. The strength (DHS signal) of mouse loci mapped from human enhancers tends to correlate with strength of the human enhancers, but a fraction of the identified orthologous pairs lacks such conservation of activity due to sequence divergence (**Fig 1G**). To rank predictions, we explored different ways to predict functional conservation between orthologous human and mouse enhancer pairs identified by *gkm-align* (**Fig 4E**). We observed that the DNase signal of a query human sequence (qDNase) was correlated with the DNase signal of the mapped mouse ortholog (mDNase) with median correlation of 0.29 (min: 0.13; max: 0.46), and the product of qDNase and gkm-similarity between query human enhancer (HE) and the mapped mouse ortholog (ME), lead to increased median correlation of 0.41 (min: 0.23; max: 0.60) (**Fig 4F**). When this product is further multiplied by enhancer prediction of ME by human trained gkm-SVM (0 ≤ *P*_*H*_(*ME*) ≤ 1), median correlation further improves to 0.46 (min: 0.25; max: 0.61). This triple product has a high value for a mapped mouse ortholog if it has similar gapped-kmer composition, a strong human enhancer ortholog, and contains conserved TFBS motifs. Training a regression model with these combinatorial features leads to median correlation of 0.55 (min: 0.32; max: 0.68) (**Fig 4F**; **Methods**). All of our human/mouse conserved enhancer mappings are ranked with these gapped-kmer based conservation scores, which we believe will facilitate downstream experimental testing by providing confidence ranking scores for functional conservation in mice.

Many of the novel enhancers identified by *gkm-align* are supported by additional evidence of conserved function. For example, gkm-align predicts a conserved enhancer in OTU Deubiquitinase 7A (OTUD7A) which is highly expressed in both human and mouse brains^55^ (**Supp Fig 13**) and is associated with a wide range of neurological diseases as such schizophrenia and epilepsy^56^. OTUD7A knockout leads to morphological deformation of cortical neurons and frequent seizure-like events in mice^57,58^. We identified orthologous pairs of putative OTUD7A enhancers in intron 1 of human and mouse OTUD7A (**Fig 5A-B**; enhancers: yellow-highlighted; hg38/chr15:31740273-31740573; mm10/chr7:63554547-63554847). Both human and mouse elements exhibit strong DNase-I accessibility and H3K27ac histone modification across biological samples related to the nervous system (**Fig 5A-B).** The two enhancers appear to have three clusters of conserved DNA base pairs with high local conservation in gapped-kmer composition (local conservation rate represented as the logo heights in **Fig 5C**), and one of the clusters located at the centers of the human and mouse enhancers contains a NEUROG/ATOH1 binding motif (GCAGATGG), which is identified among the top brain enhancer gapped-kmer weights for both human and mouse as shown in **Fig 1C**. This region has largest delta-SVM^24^ score for gkm-SVM models trained on both human and mouse brains (**Fig 5C**). Despite the clear conserved biochemical signatures and binding motif, this enhancer is predicted to be deleted in mouse by LASTZ/LiftOver (minmatch 0.01).

To show further examples of the top conserved enhancers in **Fig 5D**, we ranked enhancers that have the strongest combined (i) DNaseI accessibility, (ii) gkm-similarity, and (iii) interspecies gkm-SVM prediction, using the regression score described in **Fig 4E**. This combined regression score increases the likelihood of functional conservation as shown in **Fig 3H** for HBB CRISPRi. We ranked enhancers collected from five diverse human and mouse cell/tissues (brain, kidney, stomach, muscle, B-cell) from among the top 1% conserved enhancers with highest regression score. Among these top orthologous enhancers, we selected a subset of enhancers in the vicinity of orthologous genes with cell/tissue specific expression^55^ (**Fig 5D**; **Supp Fig 13**). These genes include KCND3 (brain; voltage-gated potassium channel subunit), PAX2 (kidney; TF associated with renal malformation^59^), GATA6 (stomach; definite endoderm TF^14^), MYOD1(muscle; TF associated with myopathy^60^), and IKZF3 (B-cell; TF mutated in leukemia^61,62^). For each of the 45 cell/tissue pairs, we generated a table of ranked orthologous human-to-mouse enhancer pairs. In addition to providing an expanded catalog of conserved distal enhancer elements, the ranking can be used to prioritize elements for functional characterization.

## Discussion

Model organism studies have elucidated many of the mechanisms of transcriptional regulation and have functionally validated the roles of some enhancers associated with human diseases. However, characterization of enhancers through model animals is possible only for enhancers with identifiable orthologs in model animals. Mice are perhaps the most facile model system for human disease, but their evolutionary distance poses more of a challenge than detecting orthologous regulatory sequence in primates (**Fig 1F-G**). Past studies using conventional genome alignment and mapping algorithms (LASTZ/LiftOver) have shown that mapping orthologous enhancers with conserved regulatory activities is much more difficult than mapping promoters. We addressed whether the difficulty of enhancer mapping is a result of rapid enhancer evolution or of limitations in conventional genome alignment algorithms. To comprehensively quantify enhancer conservation, we used gkm-SVM to generate unbiased sets of enhancers for each pair of 45 human and mouse cell/tissues matched by core TF regulators (**Supplementary Table 4**). Interestingly, enhancers appear to have highly variable levels of conservation across different cell/tissue types. We show that the conservation rate of embryonic brain enhancers are about three times higher than that of adult liver, although enhancer vocabularies are highly conserved for both tissues (**Fig 2C**). The tissue variability in enhancer conservation rate is confirmed in alignment-free analyses (**Fig 2D-G**). Part of the explanation of this apparent paradox is that tissues with low enhancer conservation also have more species-specific enhancer duplication. A significant proportion of enhancer duplications appear to have resulted from transposable elements (TE), especially LTRs, and most TE-associated enhancers are species-specific (**Fig2 H-I**), partly explaining the relative lack of enhancer conservation in cell/tissues with high TE activity, such as the liver. The cell/tissue variability in enhancer conservation aligns with previously reported cell/tissue variable conservation of gene expression, which showed rapid transcriptomic divergence of the liver relative to the brain^43^. Rapid liver enhancer evolution may explain the apparent transcriptomic differences between human and mouse livers^41,42^, and may further explain the previously reported transcriptional divergence of many cell/tissue specific genes^63^. Intriguingly, in cell/tissues with low enhancer conservation, enhancers with no sequence overlap with repetitive elements also showed reduced conservation (**Supp Fig 9B**). Our observation is consistent with a model of evolution in which TEs provide influx of novel TF binding sites through transposition and further facilitate turnover of nearby enhancers by supplying functional redundancy^64^. TEs can contribute up to 30% of the strongest DHS peaks in some tissues, but we do not know what proportion of these TFBS-carrying TEs act as transcriptional enhancers to relevant genes, and which might be regulatory noise. Only a small proportion of TE-enhancers have so far been functionally tested^48^, but we expect to see more functional validation of these elements in near future due to advances in noncoding CRISPR-based screening methodologies^54,65^.

We developed a gapped-kmer based novel alignment algorithm to detect conserved enhancers, *gkm-align. gkm-align* maps orthologous enhancers by finding alignment paths of maximal gapped-kmer composition at the resolution of sliding ∼300 base pair windows. We used a whole genome alignment strategy and present a new set of conserved enhancer predictions for human and mouse. We evaluate these predictions on 45 pairs of matched tissues using ENCODE data and show that *gkm-align* detects thousands of conserved enhancers missed by conventional alignment methods (**Figure 3I**). We further extend these predictions by combining tissue specific TF information, which predicts an additional 500 enhancers per tissue on average, and up to an 80% increase in some tissues (**Figure 4B-D**). While our analysis confirms that mapping orthologous enhancers between distant mammals is an inherently difficult problem due to rapid enhancer evolution, we show that we detect conserved enhancers of biomedical significance missed by LiftOver/LASTZ, including an intronic enhancer of OTUD7A which is associated with epilepsy in humans and when knocked out reduces dendritic density and promotes seizures in mouse. Despite the algorithmic improvement for mapping orthologous enhancers, our analysis confirms the overall weak enhancer conservation relative to promoters and that enhancers have surprisingly variable conservation rate across cells/tissues. Lastly, we provide an expanded catalogue of orthologous human-mouse enhancers, each annotated with predictive gapped-kmer based functional conservation scores. We expect that this expanded and quantitatively ranked catalogue of conserved enhancers will facilitate discovery and functional characterization by prioritizing enhancers for testing in model animals (**Fig 4E-F**, **Fig 5**).

## Supporting information

supplemental algorithm details

## Methods

### Generating enhancer and promoter sets from DNase-seq data and gkm-SVM training

For all the ENCODE DNase-seq data used in this study, DNase-seq filtered alignment bam files were downloaded from the ENCODE portal https://www.encodeproject.org/. For the primate fibroblast DNase-seq data, primate fibroblast DNase-seq raw fastq files (under GEO accession GSE129034) were downloaded from (Edsall 2019)^1^ and mapped to chimpanzee (panTro6), gorilla (gorGor5), orangutan (ponAbe3), and rhesus (rheMac8) genomes using bowtie2^2^ (-L 20). DHS peaks were called by running MACS2^3^ on bams from combined replicates using default parameters but more stringent p-value (-p 1e-9). 300 bp peaks were generated by extending +/- 150bp from MACS2 summits.

Promoter sets were generated by selecting all peaks within 2kb of any annotated TSS. Remaining distal cell specific peaks were further filtered by removing all peaks called in more than 30% of ENCODE DHS samples (hg38ubiq30.bed and mm10ubiq30.bed). This definition of promoters and enhancers was used to train the promoter and enhancer gkm-SVM models that are publicly available on the ENCODE portal (**Supplementary Table 2-3** for ENCODE accession IDs). To define primate enhancers, coordinates in hg38ubiq30.bed were mapped to the respective primate genomes by LiftOver. For **Figure 1-2**, more stringent definition was used for defining promoter elements to filter out proximal DHSs that are accessible in less than 30% of DHS samples.

For gkm-SVM model generation, gkm-SVM models were trained on top 10,000 300bp enhancers or top 10,000 300bp promoter peaks (with highest DNase-seq MACS2 peak score)vs GC and repeat matched random sequence following (Beer Shigaki Huangfu 2020; Ghandi 2014; Lee 2015)^4–6^ using default parameters (-l 11 -k 7).

To generate gkm-SVM kmer-weight vectors, A fasta file containing all 11-mers was generated using nrkmers.py from the lsgkm software package (v0.1.1) package^7^. Kmer weight vector for each 1,270 human and 153 mouse gkm-SVM enhancer models were generated using the lsgkm package’s *gkmpredict* on the 11-mer fasta file using respective gkm-SVM models.

### Interspecies cis-regulatory element prediction

Top 10,000 elements for each DHS class (enhancer, promoter) in human and mouse were used as *positive* set, and randomly sampled 300bp genomic loci with matched GC-content and repeat annotations were used as *negative set*. Gkm-SVM model trained in one species was used to make prediction for elements in each *positive* and *negative* set in the other species. AUROCs were computed from these interspecies prediction values.

### Generation of 45 human-mouse cell/tissue pairs

For each human sample (N = 1,270; **Supplementary Table 2**), we identified its best matching mouse sample (N = 153; **Supplementary Table 3**) by finding mouse sample with the highest gkm-SVM kmer weight Pearson correlation. The best matching human sample for each mouse sample was identified similarly. The 45 human-mouse cell/tissue pairs were derived as those human and mouse samples that reciprocally mapped to each other by top matched kmer weights (**Supplementary table 4**).

### Visualization of epigenetic signals at DHSs

We used deepTools’s computeMatrix (v3.5.1)^8^ to compute average epigenetic signals across all DHS for each -1000 to 1000 base pair positions (resolution: 10) relative to DHS center using the following command:

computeMatrix reference-point -S example.bigWig -R example.bed -a 1000 -b 1000 --referencePoint center -bs 10 -o example.tab.gz

ENCODE ChIP-seq and DNase-seq bigwig files used for this analysis (**Fig 1B**, **Supp Fig 1-2**) are listed in **Supp Table 1**.

### Defining orthologous syntenic intergenic loci in human and mouse

The list of all human-mouse orthologous protein-coding gene pairs (N = 15,712) was obtained from the mouse ENCODE consortium publication^9^ (**Supplementary Table 5**). 15,500 out of 15,712 gene pairs with their human and mouse gene ID’s also present in the Ensembl database were used, and the remaining 212 genes were filtered out (Homo_sapiens.GRCh38.96.chr.gtf, Mus_musculus.GRCm38.96.chr.gtf). Coordinates of these conserved human and mouse 15,500 genes were extracted from the Ensembl gtf files.

To identify all human-mouse syntenic intergenic loci, we first identified all neighboring pairs of human protein-coding genes conserved in mouse. Denote such human gene pair as HG_1_ and HG_2_. If their mouse gene orthologs, MG_1_ and MG_2_, are also neighbors in the mouse genome and if the relative transcriptional directions of [HG_1_ and HG_2_] and [MG_1_ and MG_2_] are also preserved (i.e., HG_1_/HG_2_ and MG_1_/MG_2_ both have tandem, convergent, or divergent transcriptional directions), we label [HG_1_/HG_2_, MG_1_/MG_2_] as human-mouse syntenic neighboring gene pairs (N = 12,455). Human *syntenic intergenic locus* was defined as the union of genomic space between a pair of human syntenic neighboring genes and their gene bodies, and mouse *syntenic intergenic locus* was defined similarly. This led to 12,455 pairs of human and mouse genomic regions that we call “human-mouse syntenic intergenic loci.”

### Using LASTZ/LiftOver for estimating conservation rate of cis-regulatory DNA elements

LASTZ (v1.03.66) chain files for human-mouse genome alignment were downloaded from these links (June 2020):

https://hgdownload.soe.ucsc.edu/goldenPath/hg38/vsMm10/hg38.mm10.all.chain.gz

https://hgdownload.soe.ucsc.edu/goldenPath/mm10/vsHg38/mm10.hg38.all.chain.gz

LiftOver software was downloaded from this link (June 2020): https://hgdownload.soe.ucsc.edu/admin/exe/linux.x86_64/liftOver

Minmatch=0.01 and -multiple options were used for high sensitivity and to allow duplicate mappings. Command to map human elements to mouse:

liftOver h_elems.bed hg38.mm10.all.chain.gz h_elems_mapped.bed h_elems_not_mapped.bed -minMatch=0.01 -multiple

Command to map mouse elements to human:

liftOver m_elems.bed mm10.hg38.all.chain.gz m_elems_mapped.bed m_elems_not_mapped.bed -minMatch=0.01 -multiple

With these settings, LASTZ/LiftOver maps a human DHS (e.g., human brain enhancer) to 1≥ mouse locus, and we say it is *conserved* if at least one of the mouse mapped loci overlaps (≥1 bp) with a mouse DHS in the matched cell/tissue in the matched syntenic intergenic loci. Human DHS conservation rate is defined as the fraction of conserved human elements, and mouse DHS conservation rate is computed similarly. Human-mouse conservation rate is defined as the average of the two rates.

### Sequence homology analysis for quantifying the proportions of orthologous and paralogous enhancers

Sequence similarity between a pair of enhancers was quantified as the similarity in gapped-kmer composition (*gkm-similarity*; defined in main **Fig 3A** and **Appendix B**) using gkmSVM^5^ R software package (v0.81.0). Prior to computing gkm-similarities, we masked portions of enhancer sequences that are predicted to be highly prevalent genome-wide to prevent trivial sequence matches by ubiquitous sequence patterns such as the low-complexity repeats (**Supplementary information E.3**; **Supp Fig 8**). About 10% of enhancer sequence base pairs were masked and replaced with random base pairs.

To compute proportions of human orthologous and paralogous enhancers in each human/mouse cell/tissue pairs (e.g., human brain and mouse brain), we obtained top 5,000 enhancers with highest DNase I signals (MACS2 peak score) and computed their pairwise sequence similarity with all other 5,000 human enhancers and with every mouse enhancer of the matched cell/tissue type. If the number of mouse enhancers exceed 50,000, we used the top 50,000. Using these values, we identified top matched human and mouse enhancers that are most similar to each of the top 5,000 human enhancers. Sequence similarities with top matched human and mouse enhancers were then used to classify a human enhancer as orthologous, paralogous or neither. Denoting top sequence similarity with human enhancers and with mouse enhancers as y and x respectively, we classified human enhancers according to the following classification rules:

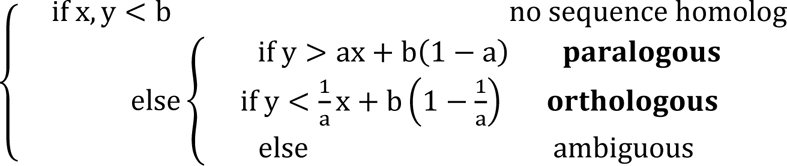

, where (a, b) = (4, 0.1). The decision regions for orthologous and paralogous enhancers are each shaded blue and red in **Figure 2E**. Mouse enhancers were classified similarly.

### Annotations for repetitive DNA elements

Annotations for repetitive elements were downloaded from the following links^10^:

http://hgdownload.cse.ucsc.edu/goldenpath/hg38/database/rmsk.txt.gz

http://hgdownload.cse.ucsc.edu/goldenpath/mm10/database/rmsk.txt.gz

### Quantifying enhancer strength of human loci mapped from mouse HBB enhancers using CRISPRi perturbation data

HCR-FlowFISH CRISPRi data (K562 cell line; HBE1 expression perturbation as readout) were downloaded from the ENCODE portal (accession ID provided in **Supplementary Table 1**)^11,12^. Two biological replicates were used, where each replicate generates sgRNA sequence read counts for low and high expression sort bins. The following equation was used to compute CRISPRi effect size of each sgRNA.

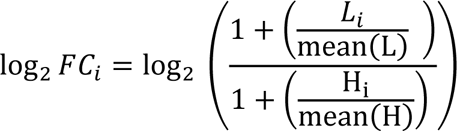

where L and H are each a vector encoding the number of reads for each sgRNA in low and high sort bins respectively. Normalization with mean underweight sgRNAs with low read counts^12^. Enhancer strength of putative human enhancers, mapped from mouse HBB enhancers using gkm-align, were computed as average log2FC of sgRNA target that overlap with each enhancer. Mouse enhancer coordinates were defined using mouse embryonic liver DHS (ENCFF578VRG; 300 base pair wide extended from the summit) that also overlap with GATA1 ChIP-seq peaks in mouse erythroblast (ENCFF676TDJ) within mm10/chr7:103851395-103883181.

### Regression model for predicting functional conservation

For predicting functional conservation of human enhancers in mouse (measured as DNase-signal at mouse loci mapped from human enhancers), we denote a set of human enhancers as {*HE*_1_, *HE*_2_, … } and denote mouse elements mapped from query *HE* by gkm-align as *ME*_*i*_. Denoting DNase-signal at query *HE* as *q*_*sig*_and signal at mapped *ME* as *m*_*sig*_ (fold change relative to genomic average), we model functional conservation as:

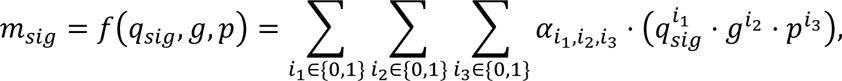

where *g* = *gkmsim*(*HE*_*i*_, *ME*_*i*_), *p* = *P*_*H*_(*ME*_*i*_), and *f* linearly combines all multiplicative combinations of the three variables (**Fig 4E**). Predicted mouse regulatory signal 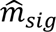 is computed using 5-fold CV linear regression. *q*_*sig*_ ⋅ *g* ⋅ *p*, which also has high predictability (**Fig 4F**), may also be used instead, if *m*_*sig*_ is not accessible (e.g., no relevant functional experiments performed in mice).

## Acknowledgement

JWO and MAB were supported by NIH grants R56 HG012110, R01 HG012367, and U01 HG009380 to MAB. We thank members of the Beer lab for useful discussions.

## Supplementary Tables

A description of the contents of all Supplementary Tables is provided below.

**Supplementary Table 1**: ENCODE IDs of experiments used for each figure.

**Supplementary Table 2**: Human ENCODE IDs for DNase-seq experiments and corresponding gkm-SVM enhancer models.

**Supplementary Table 3**: Mouse ENCODE IDs for DNase-seq experiments and corresponding gkm-SVM enhancer models

**Supplementary Table 4**: List of 45 cell/tissue pairs matched by the gkm-SVM enhancer models

**Supplementary Table 5**: Human-mouse orthologous protein-coding genes from Mouse ENCODE^9^

Additional tables/files are accessible through https://beerlab.org/gkmalign/

## Supplementary information: gkm-align algorithmic details & code access

The algorithmic details of gkm-align are provided in the separate supplementary file named ‘Supplementary_information_algorithm.pdf’

gkm-align software can be downloaded from: https://github.com/oh-jinwoo94/gkm-align

Command lines to run gkm-align are listed in the github README.

## Supplementary Figures

**Supplementary Fig. 1:**
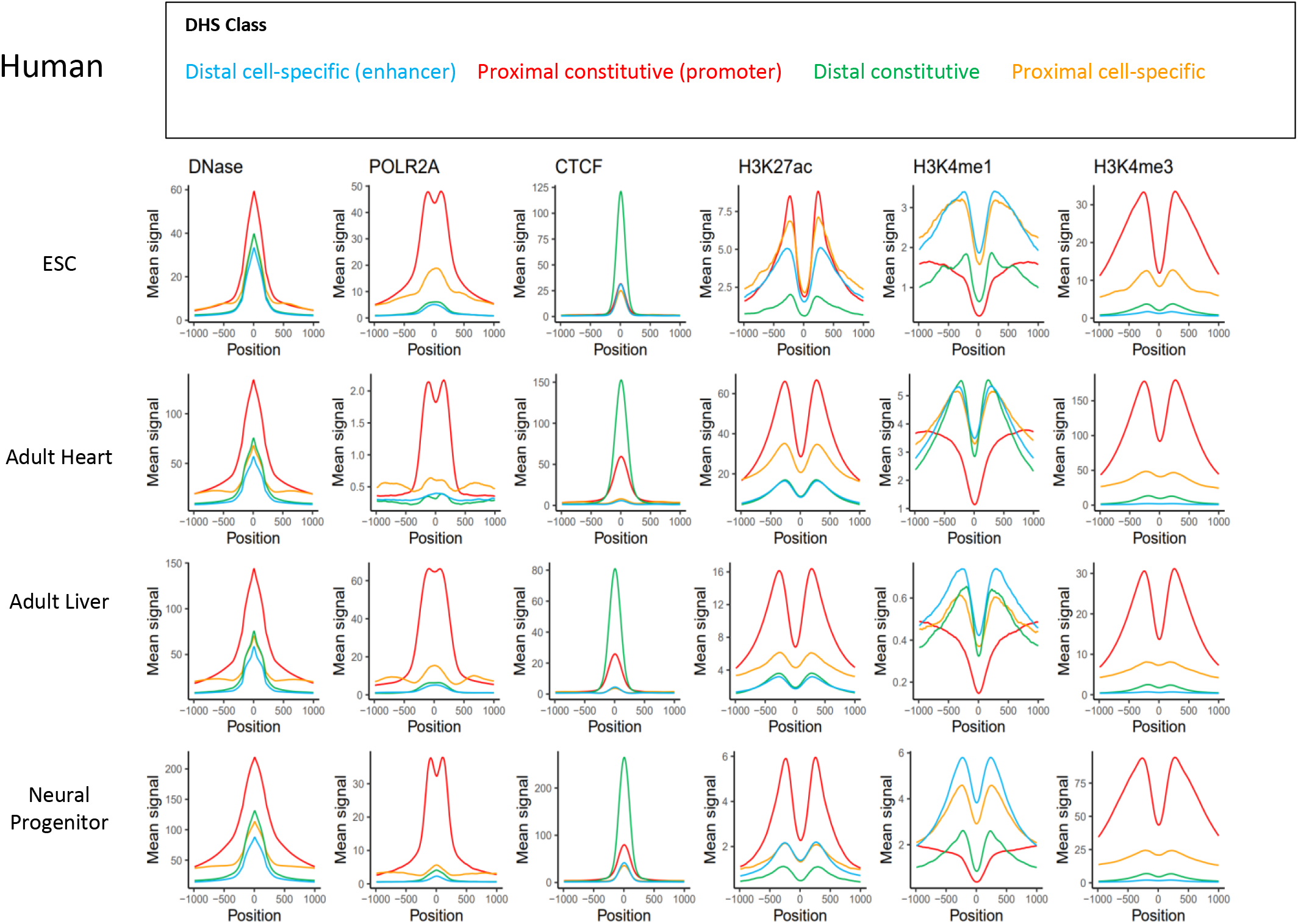
Epigenetic profiles of DHS subclasses across diverse human cell/tissues. Classification of DHSs from diverse cell/tissues by their distances to nearest TSS (proximal: < 2kb; distal: > 2kb) and cell-specificities of chromatin accessibilities (cell-specific: DHS in less than 30% of all biosamples; constitutive: otherwise) and visualization of average epigenetic signals around DHS peak centers by DHS class. ENCODE IDs of the experiments for each cell/tissues are listed in **Supp Table 1**.

**Supplementary Fig. 2:**
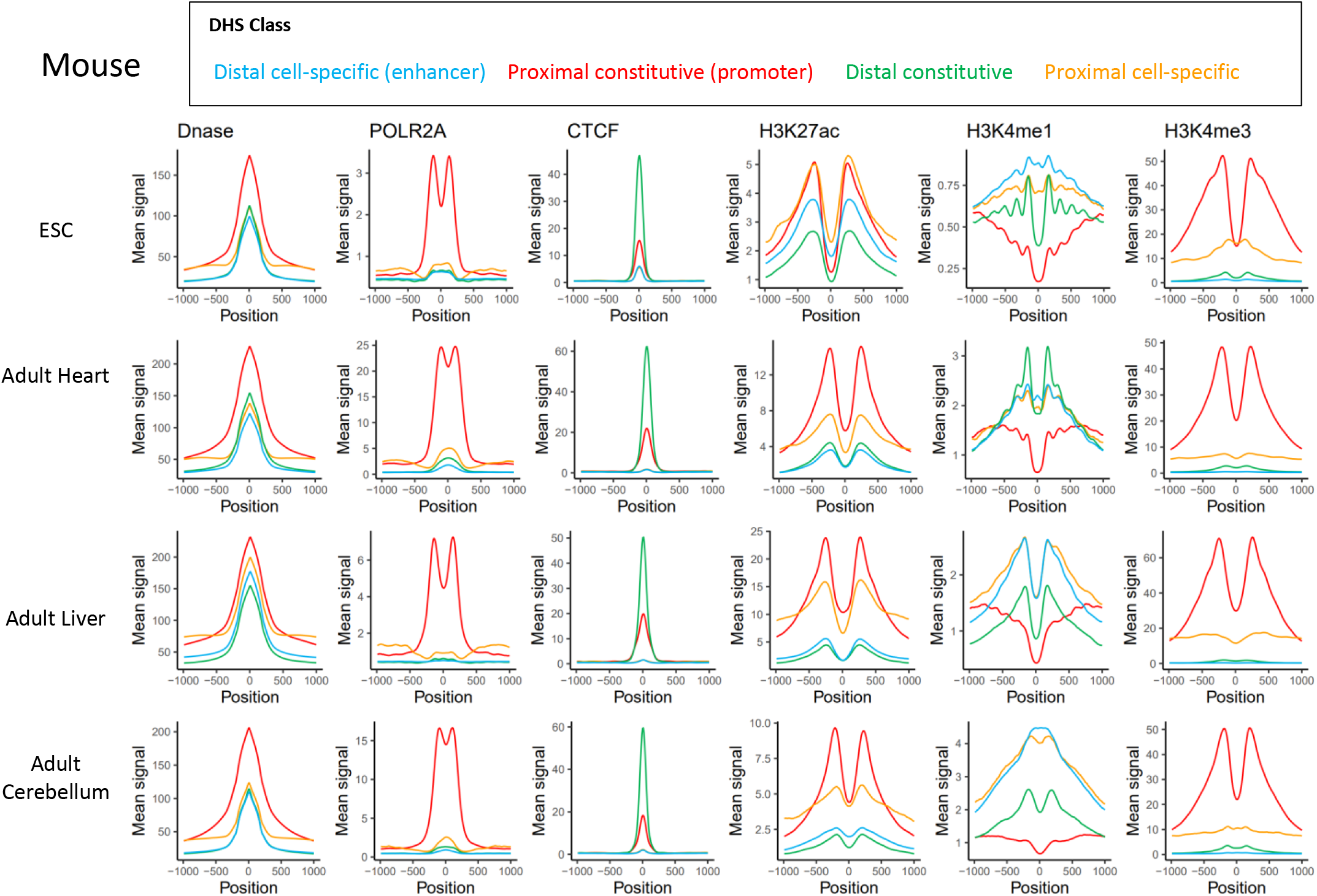
Epigenetic profiles of DHS subclasses across diverse mouse cell/tissues. Classification of DHSs from diverse cell/tissues by their distances to nearest TSS (proximal: < 2kb; distal: > 2kb) and cell-specificities of chromatin accessibilities (cell-specific: DHS in less than 30% of all biosamples; constitutive: otherwise) and visualization of average epigenetic signals around DHS peak centers by DHS class. ENCODE IDs of the experiments for each cell/tissue are listed in **Supp Table 1**.

**Supplementary Fig. 3:**
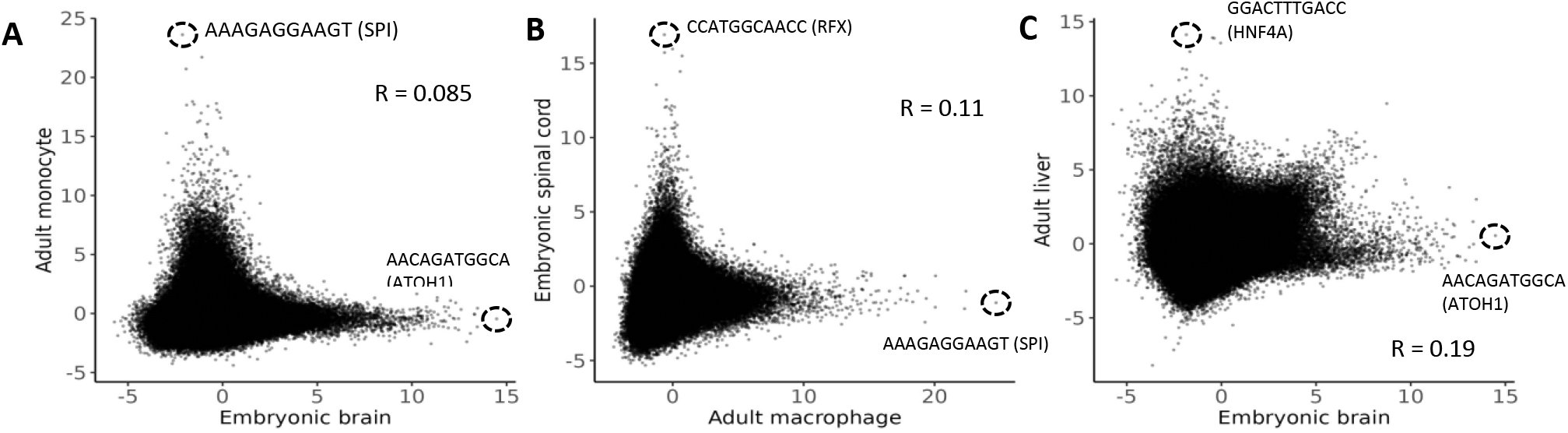
gkm-SVM enhancer vocabularies are distinct for distinct cell/tissue types. Comparing kmer-weights between **A)** monocyte & brain **B)** spinal cord & macrophage **C)** brain & liver. Gkm-SVM models used for this figure can be identified by their model aliases and searched in the ENCODE portal (brain = DHS_399_hg38; spinal cord = DHS_421_hg38; monocyte = DHS_828_hg38; macrophage = DHS_482_hg38; liver = DHS_1229_hg38). More detailed information about these models is given in **Supplementary Table 2**.

**Supplementary Fig. 4:**
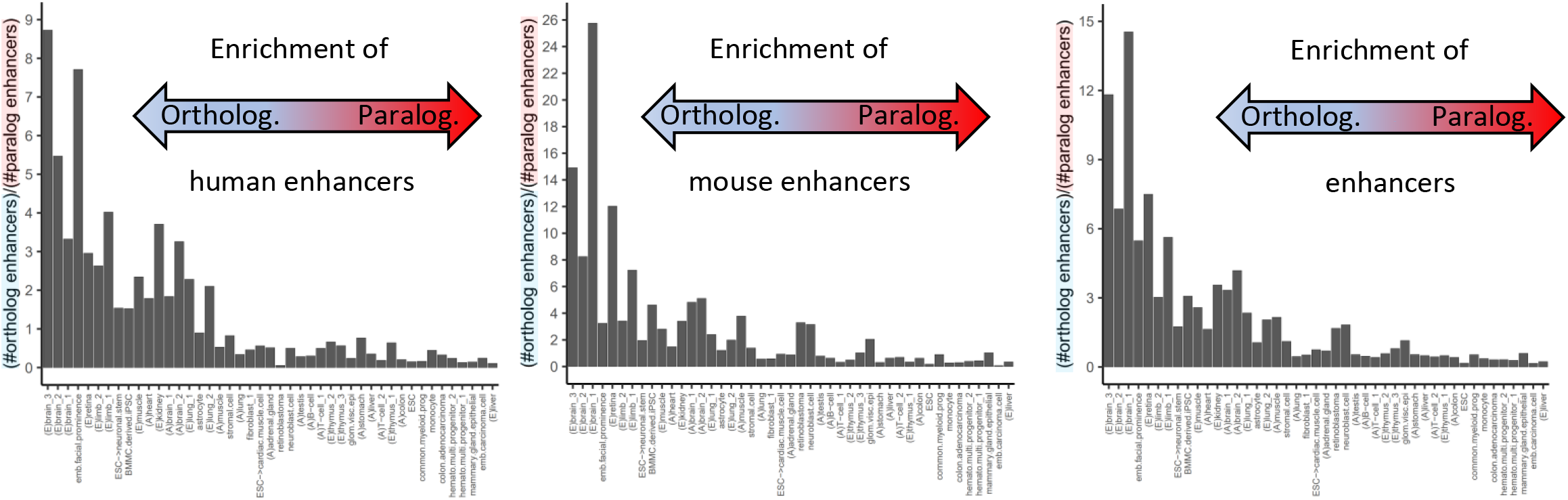
Ratio of orthologous to paralogous enhancers across the 45 cell/tissue pairs. Ratio of orthologous to paralogous enhancers for (A) human (B) mouse (C) and their average (**Methods**). Cell/tissues are listed in the same order as **Fig 2A**.

**Supplementary Fig. 5:**
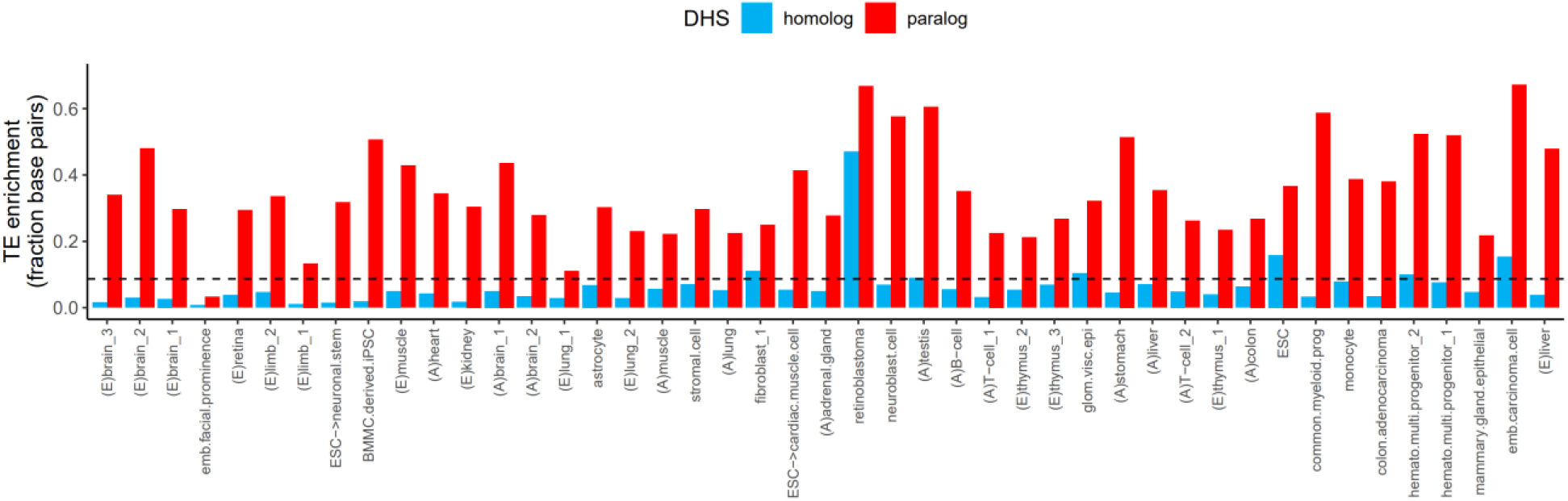
LTR transposable elements are enriched in paralogous enhancers. Fraction of base pairs annotated as LTR transposable element for human enhancers classified as homologous or paralogous enhancers. Dotted line indicates the fraction of human genome annotated as LTR transposable element (0.087).

**Supplementary Fig. 6:**
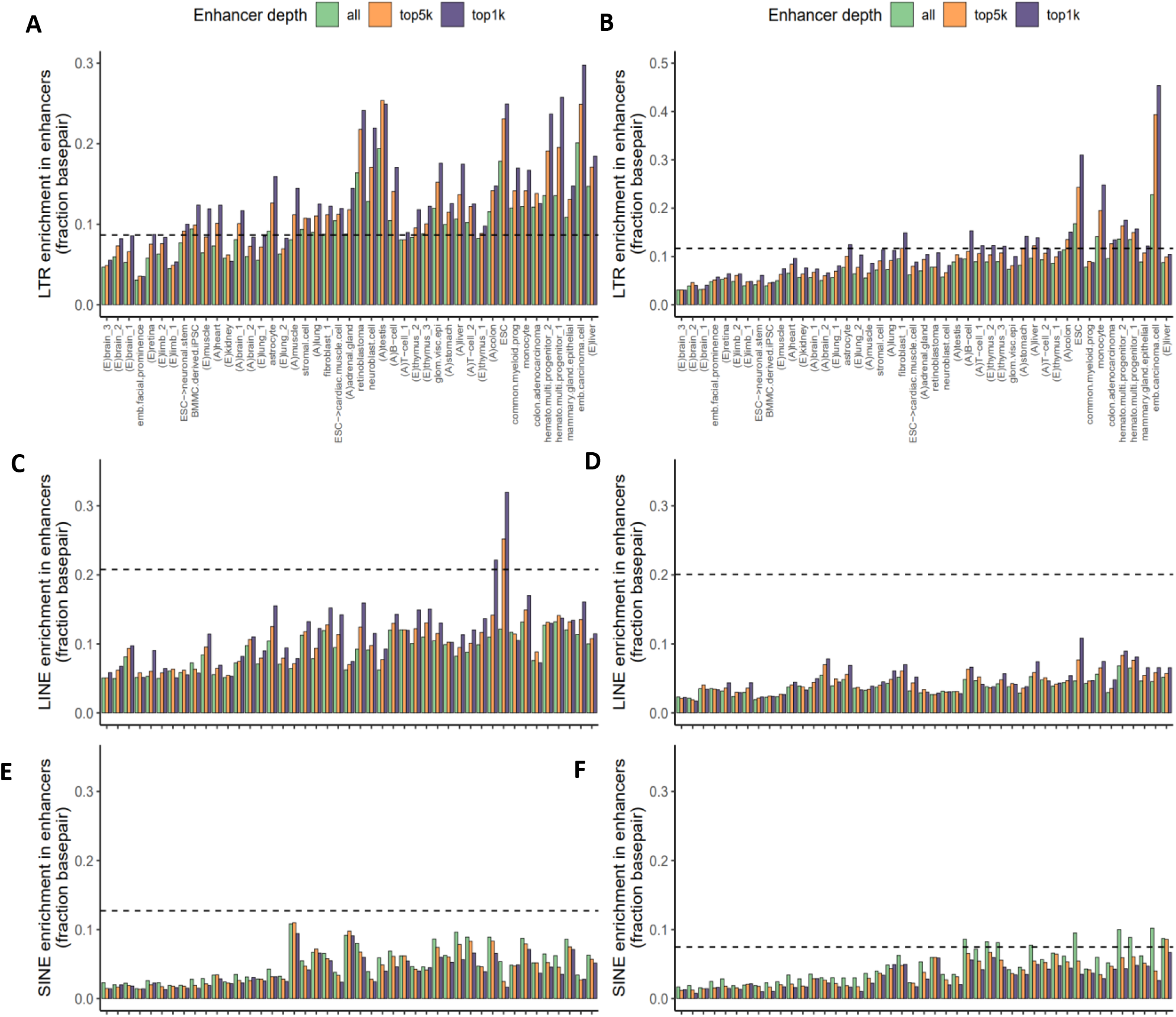
Enrichment of transposable element in enhancers by enhancer signal depths. Fraction of enhancer base pairs that overlap with LTR (**A, B**), LINE (**C**, **D**) and SINE (**E**,**F**) transposable elements for human (**A**, **C**, **E**) and mouse (**B**, **D**, **F**) across the 45 cell/tissues. The bar plots also show TE-enrichment for top 5,000 and 1,000 enhancers with highest DNase signal in each cell/tissue pairs. Dotted line indicates the fractions of human or mouse genomes covered by LTR annotation (human: 0.087 [**A**], mouse: 0.12 [**B**]), LINE (human: 0.21 [**C**], mouse: 0.20 [**D**]), and SINE (human: 0.13 [**E**], mouse: 0.075 [**F**]) transposable elements. Identical tissue orders for all **Supp Fig 6A-F** and **Fig 2A**.

**Supplementary Fig. 7:**
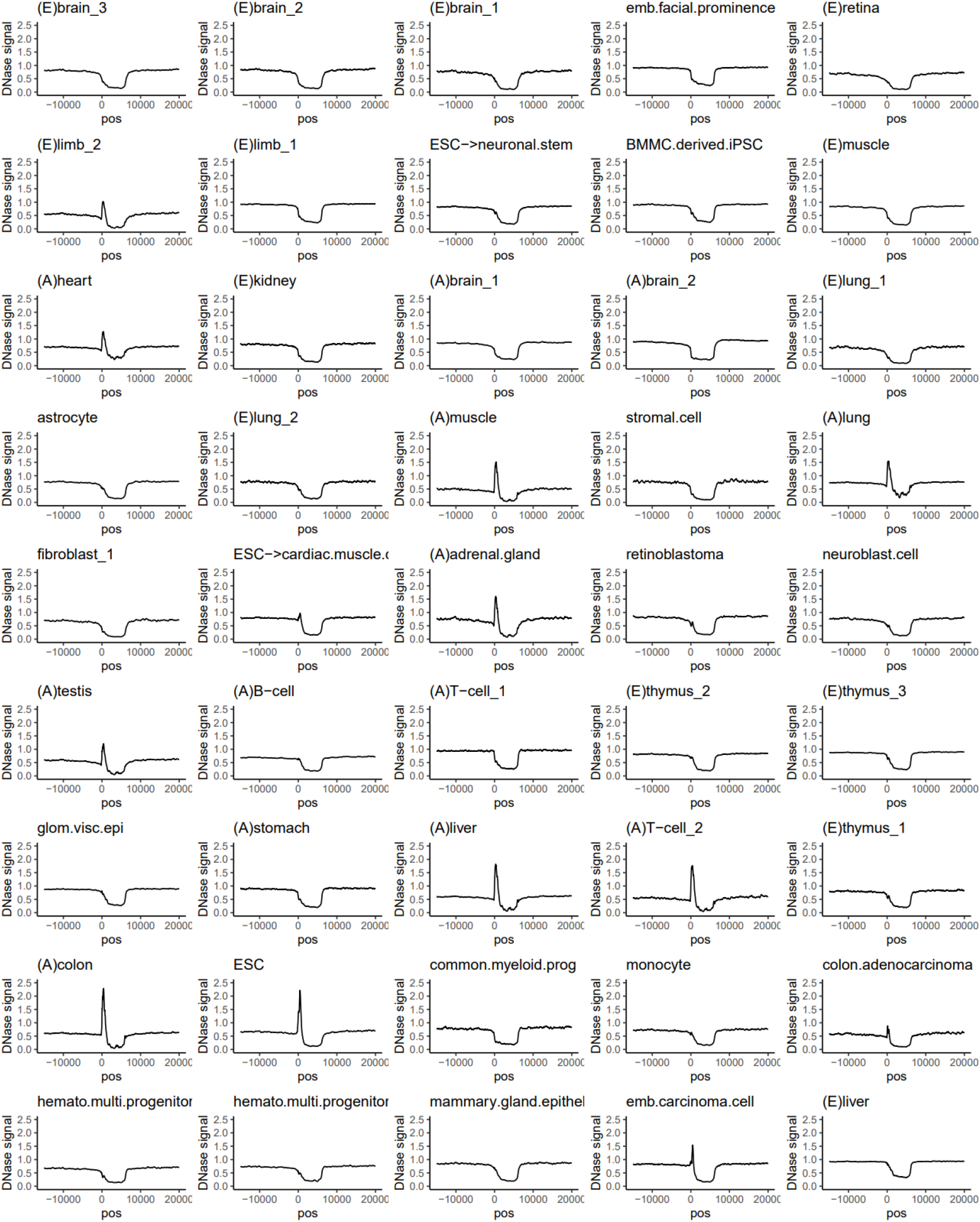
Average DNase accessibility profiles of full length human LINE1 elements. Average DNase accessibility profiles of full length (5,000 < size < 7,000) human LINE1 elements across the 45 cell/tissues. Position is relative to LINE1 5’ end along their transcriptional directions. All cell/tissues show depletion of accessibility along LINE1 bodies while a subset of cell/tissues show high accessibility at 5’ end (TSS). DNase signals computed as fold changes from genomic average.

**Supplementary Fig. 8:**
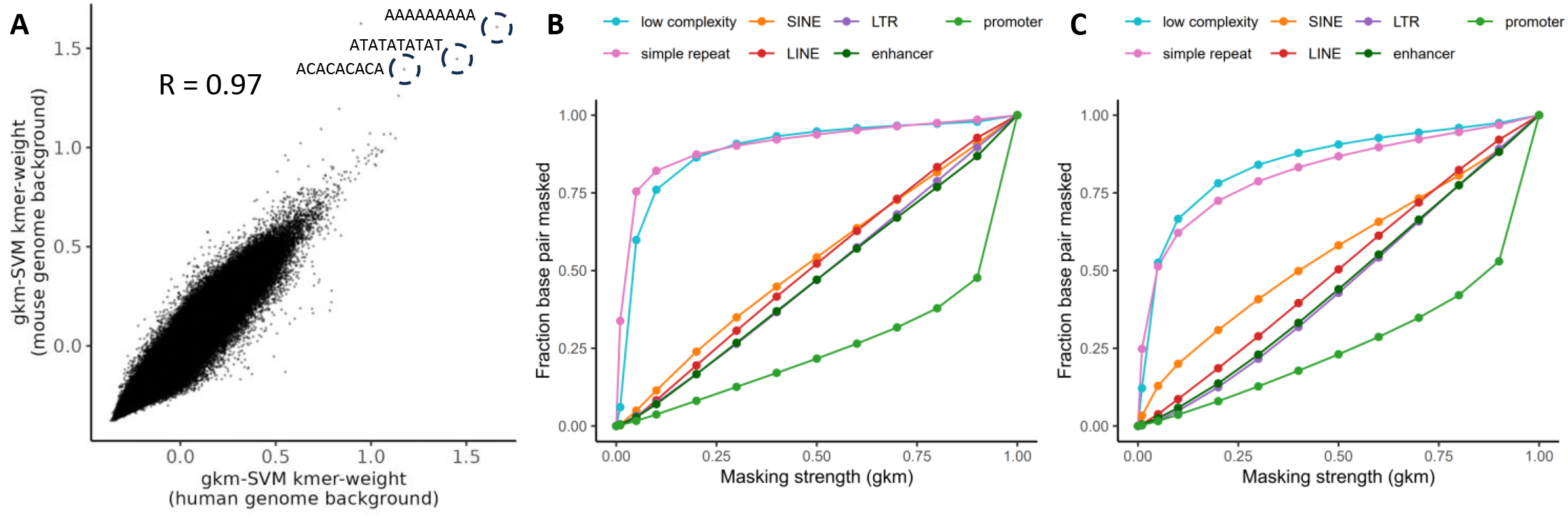
Detecting and masking repetitive DNA sequence patterns using gkm-SVM. **A)** Comparison of repetitive gkm-SVM vocabularies obtained from human (x-axis) and mouse (y-axis) by kmer weight correlation (R = 0.97). **B)** Fraction of base pairs that are masked by gkm-SVM repeat masking for different classes of repetitive elements at varying masking strengths (method described in **Supplementary information E.3**) in human and **C)** in mouse.

**Supplementary Fig. 9:**
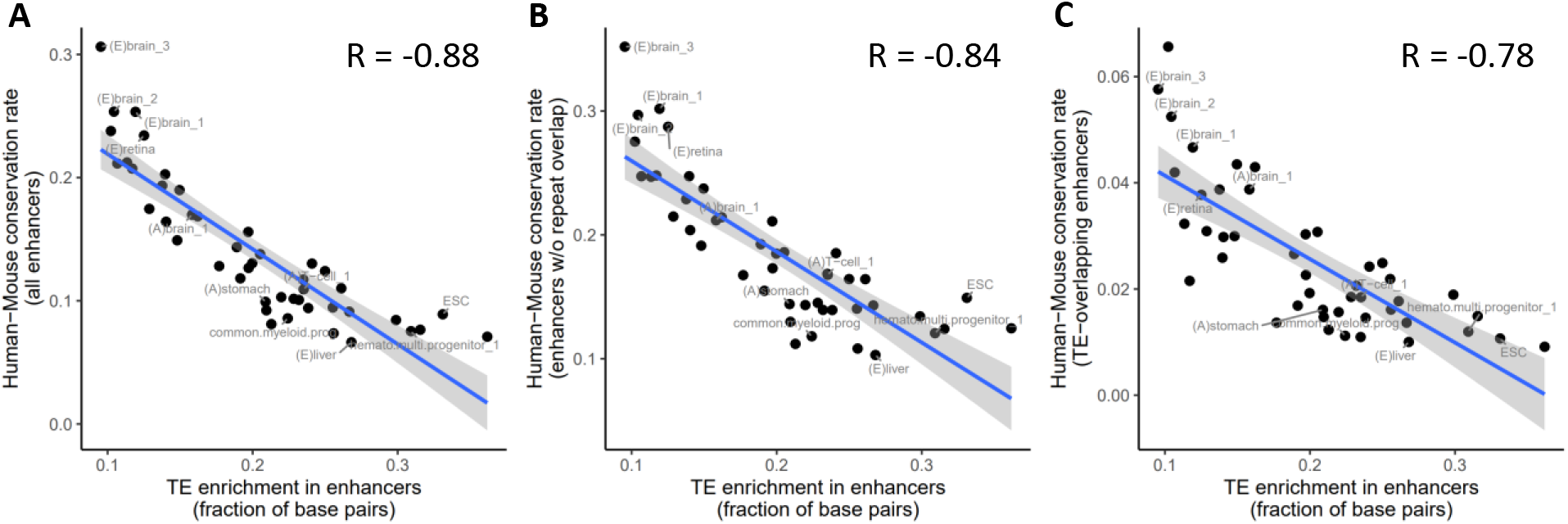
human-mouse enhancer conservation rate decreases with TE-association. **A)** Cell/tissues with high TE enrichment in enhancers have lower enhancer conservation rate (R = - 0.88), which holds true for both subsets of enhancers **B)** with no repeat annotation (R = -0.84) **C)** with more than 50% overlap with LTR, LINE or SINE annotations (R = -0.78).

**Supplementary Fig. 10:**
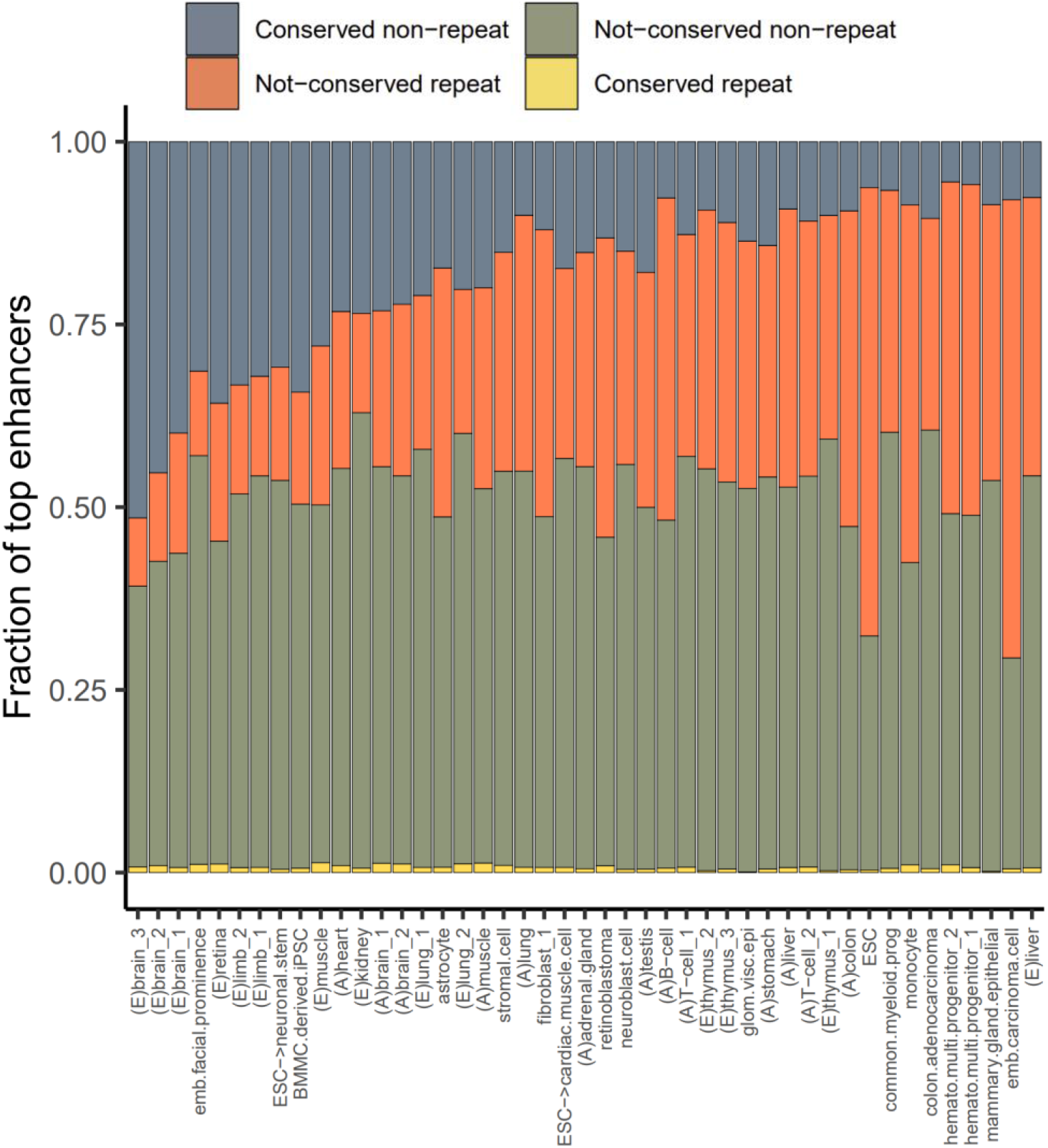
Conservation rate of strong enhancers by association with transposable elements. Same as main **Fig 2I** but using top 1,000 enhancers by DNase-signal.

**Supplementary Fig. 11:**
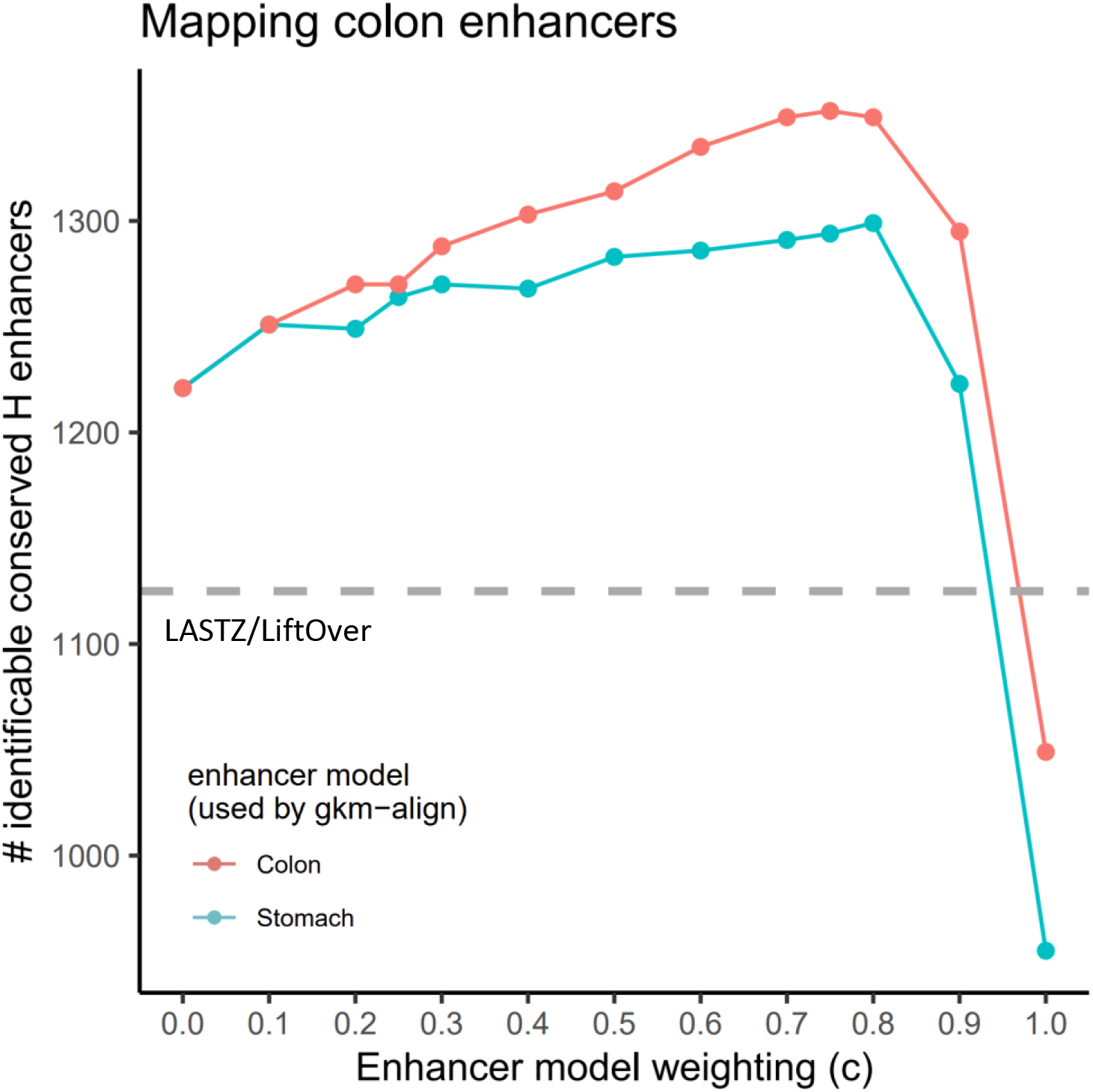
Enhancing discovery of conserved colon enhancers by incorporating gkm-SVM enhancer vocabularies. Number of human colon enhancers mappable to mouse colon enhancers using LASTZ/LiftOver (grey dashed line) and gkm-align weighted by gkm-SVM enhancer models trained on colon (red) and stomach (blue) with varying enhancer model weights.

**Supplementary Fig. 12:**
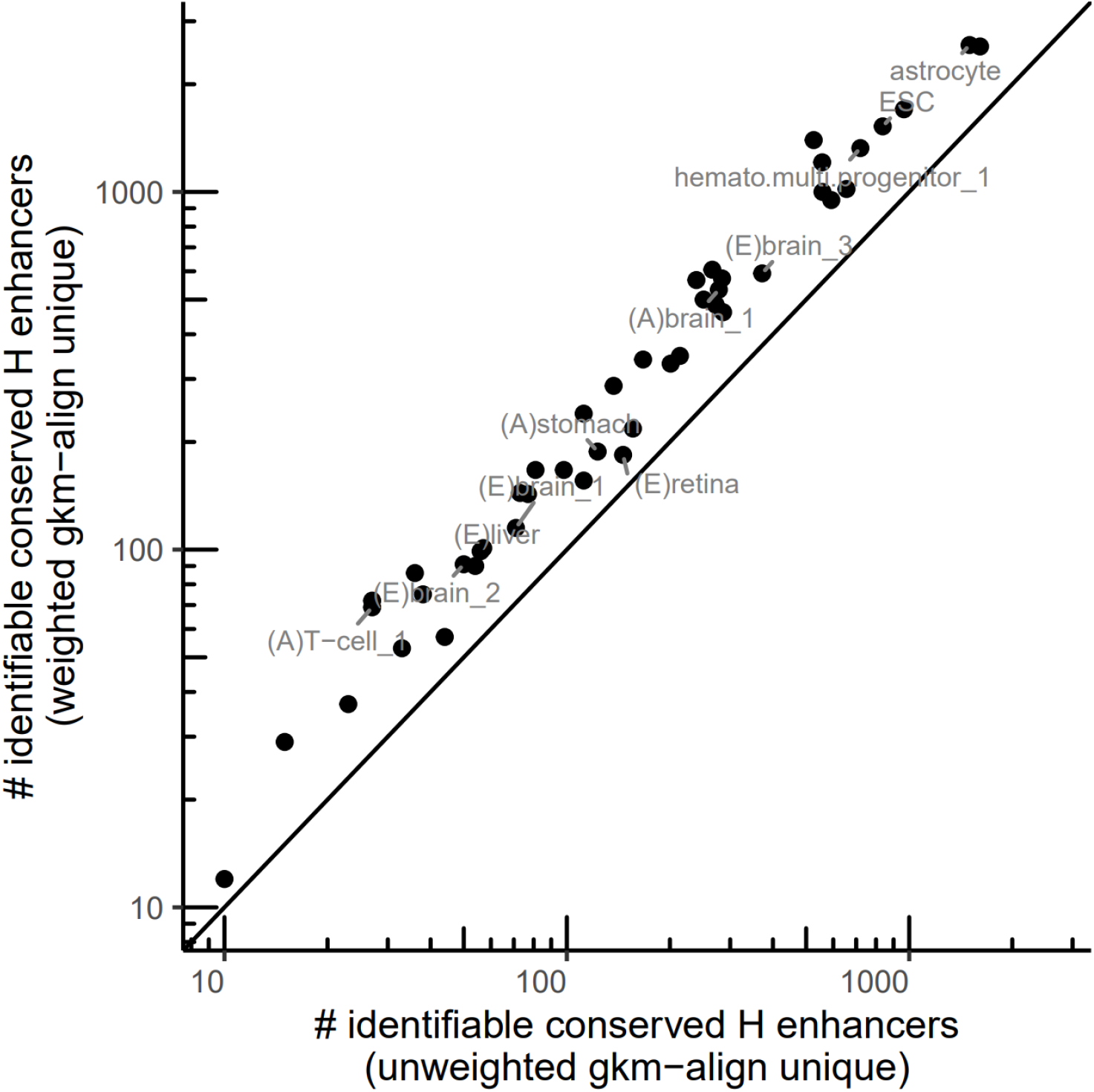
Number of unique conserved enhancer discovery by unweighted and weighted gkm-align. Number of conserved enhancers uniquely discoverable by unweighted/generic gkm-align (x-axis) or gkm-SVM weighted cell-specific gkm-align (y-axis) for each of the 45 cell/tissue pairs.

**Supplementary Fig. 13:**
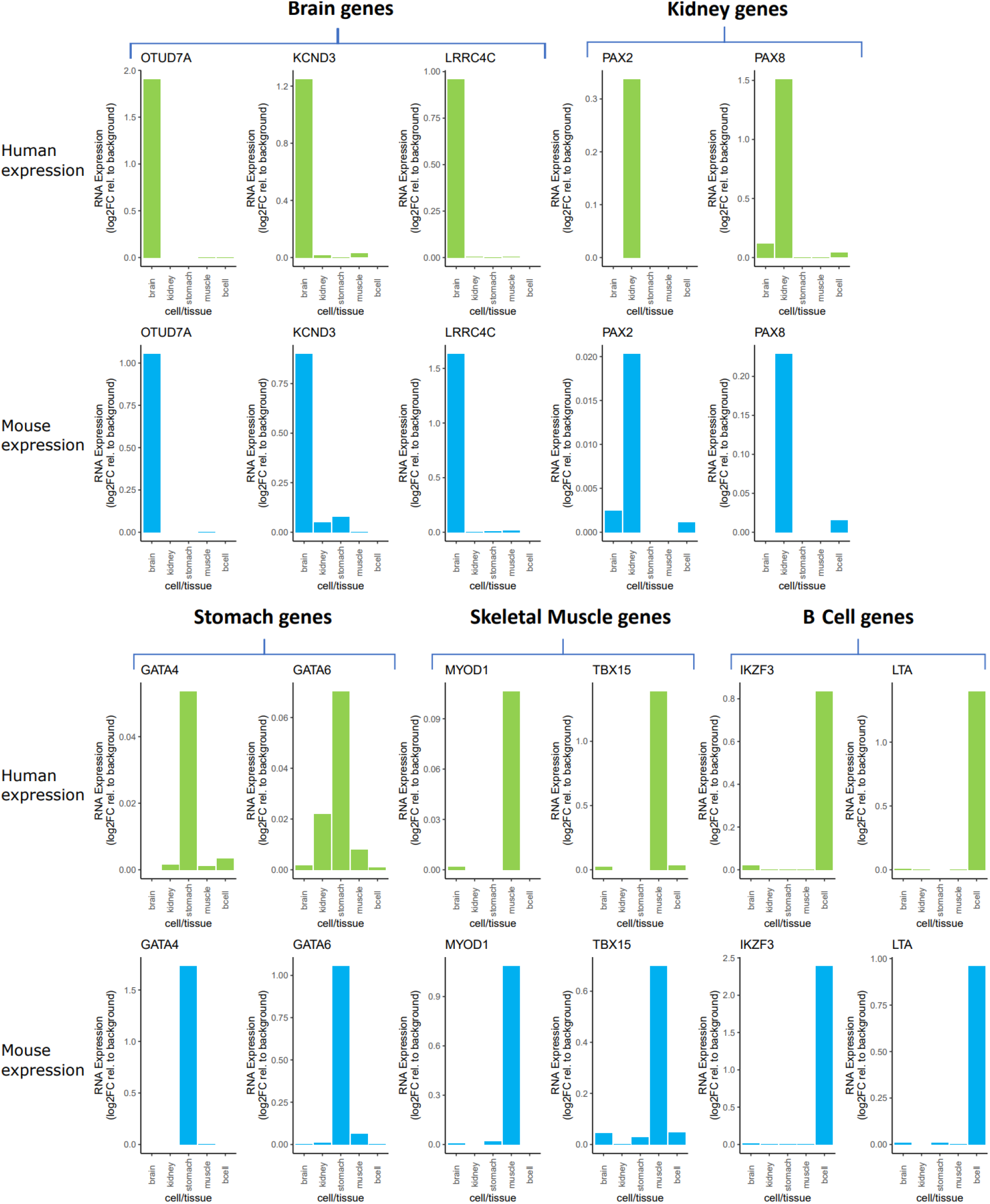
Cell/tissue specific RNA expression of genes near conserved enhancers. RNA expression level of selected cell-specific genes near conserved enhancers with top predicted functional conservation. Expression level is relative to average transcriptome-wide expression level in log2 scale (i.e. expression level of 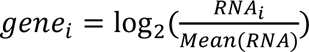). RNA-seq experiments used for this analysis are listed in **Supplementary Table 1**.

