## supplemental algorithm details for "Gapped-kmer sequence modeling robustly identifies regulatory vocabularies and distal enhancers conserved between evolutionarily distant mammals"

#### A Overview

In this document, we provide the algorithmic details of gkm-align. Machine learning models using gapped-kmers have shown that gapped-kmers can efficiently represent TF binding sites in cell-specific enhancers and regulatory vocabularies [1,2,3,4]. The gkm-align algorithm utilizes gapped-kmers as sequence feature, and identifies conserved enhancers by finding optimal alignment paths of maximum gapped-kmer similarity.

Given a human-mouse syntenic locus, gkm-align identifies an alignment path spanning the human and mouse sequences ($S_H$, $S_M$). The alignment is done at the resolution of sliding windows (**Fig.1A** [5]; e.g., window width, $w = 300$; slide step size, $s = 20$). We first compute pairwise sequence similarities (using the gapped-kmer kernel, i.e. the dot product of gapped-kmer counts) between all pairs of human and mouse sliding windows, and encode the pairwise sequence similarity values in the matrix $G$ (**Fig.1B** details in **Section B**). $G[i,j]$ encodes the gapped-kmer similarity between the $i^{th}$ and $j^{th}$ sliding windows of $S_H$ and $S_M$ (denoted as $S_{H,i}$ and $S_{M,j}$). Repetitive portions of the human and mouse sequences are masked prior to computing $G$ (**Section E**). The resulting matrix $G$ is then used to identify the optimal alignment path with maximal gapped-kmer similarity. (**Fig.1C**; details in **Section C**). Gkm-align is extended genome-wide by defining human-mouse syntenic blocks using the ENCODE human-mouse orthologous gene coordinates (**Fig.1D**; details in **section D**). Human-mouse short sequence matches are further used to obtain finer syntenic blocks, leading to higher computational efficiency of whole-genome alignment and resolution of small structural variations. Lastly, we can further improve conserved enhancer discovery of gkm-align by incorporating gkm-SVM cell-specific enhancer regulatory vocabularies (**Section E**).

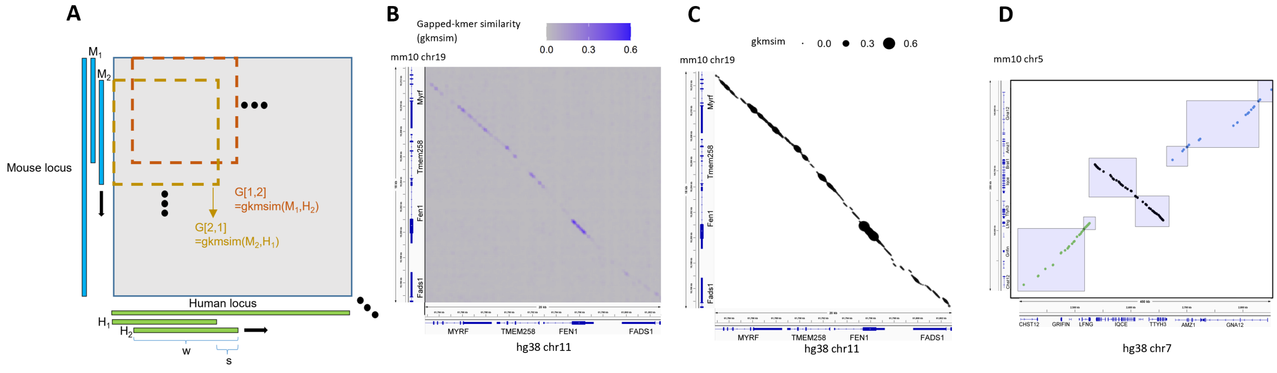


Figure 1: Graphical overview of the document

[1]Ghandi, Mahmoud, et al. "Enhanced regulatory sequence prediction using gapped k-mer features." PLoS computational biology 10.7 (2014): e1003711.
[2]Lee, D. et al. A method to predict the impact of regulatory variants from DNA sequence. Nat. Genet. 47, 955-961 (2015).
[3]Beer, M. A. Predicting enhancer activity and variant impact using gkm-SVM. Hum. Mutat. (2017).
[4]Yan, J. et al. Systematic analysis of binding of transcription factors to noncoding variants. Nature 591, 147-151 (2021).
[5]**Fig** and **Section** refer to the materials present in this current document unless noted otherwise.

### B Computing $G$: matrix encoding sequence-similarities by gapped-kmer composition

#### B.1 Comparing pairs of DNA sequences by their gapped-kmer compositions

Most of the contents in this subsection (**B.1**) were adapted from the original gkm-SVM paper. [6]

**Definition 1** *kmer(k) is a length-k sequence of* $\{A, C, G, T\}$ *.*

**Definition 2** *Gapped-kmer(l,k) is a length-l sequence of* $\{A, C, G, T, -\}$ *with k number of elements from* $\{A, C, G, T\}$.

**Definition 3** *A gapped-kmer(l,k),* $s_g$*, is contained (∈) in a kmer(l),* $s_k$*, if* $s_g[i]$ *equals* $s_k[i]$ *or* $s_g[i]$ *is a gap (-) for all* $1 \leq i \leq l$. [7] *Further,* $s_k$ *is contained (∈) in S if* $s_k$ *is a subsequence of S. Lastly, if* $s_g \in s_k$ *and* $s_k \in S$*, then* $s_g \in S$.

**Definition 4** $g_{l,k}$ *is a function that maps a sequence to its gapped-kmer composition, where* $i^{th}$ *element of* $g_{l,k}(S)$ *equals the number of* $i^{th}$ *gapped-kmer, in alphabetical order, contained in S.*

Then, gkm-similarity of $i^{th}$ and $j^{th}$ sliding windows of $S_H$ and $S_M$, denoted as $S_{H,i}$ and $S_{M,j}$, is computed as the cosine similarity of their gapped-kmer compositions:

$$\begin{aligned} G_{S_H,S_M}[i,j] &= < g(S_{H,i}), g(S_{M,j}) > \\ &= \frac{g(S_{H,i}) \cdot g(S_{M,j})}{|g(S_{H,i})||g(S_{M,j})|} \end{aligned} \quad \text{(Eq.1)}$$

Efficient computation matrix $G$ requires efficient computation of $g(S_1) \cdot g(S_2)$ for any DNA sequences $S_1$ and $S_2$. Since the number of length-l gapped kmers in $S$ can be counted by summing up the number of gapped-kmers contained in each length-l subsequences of $S$, $g(S) = \sum_{i=1}^{|S|-l+1} g(S(i,l))$. [8] Then,

$$\begin{aligned} g(S_1) \cdot g(S_2) &= (\sum_{i=1}^{|S_1|-l+1} g(S_1(i,l))) \cdot (\sum_{i=1}^{|S_2|-l+1} g(S_2(i,l))) \\ &= \sum_{i=1}^{|S_1|-l+1} \sum_{j=1}^{|S_2|-l+1} (g(S_1(i,l)) \cdot g(S_2(j,l))) \\ &= \sum_{i=1}^{|S_1|-l+1} \sum_{j=1}^{|S_2|-l+1} \begin{cases} \binom{l-m_{ij}}{k}, & l-k \geq m_{ij} \\ 0, & l-k < m_{ij} \end{cases}, \end{aligned} \quad \text{(Eq.2)}$$

where $m_{ij}$ denotes the total number of mismatched bases in a pair of length-l sequences $S_H(i,l)$ and $S_M(j,l)$ (i.e. $m_{ij} = \sum_{p=0}^{l-1}(S_H[i+p] \neq S_M[j+p])$). Gkm-align calculates $m_{ij}$ using SIMD parallel computation, which leads to the time-complexity independent of $l$.

Therefore, for any pair of sequences , $S_H$ and $S_M$, $g(S_H) \cdot g(S_M)$ is well defined as long as $|S_H|$ and $|S_M|$ are larger than or equal to $l$. This computation can further be optimized if the sliding windows overlap (as visualized in **Fig 1A**).

#### B.2 Efficient computation of matrix $G$

$G$ encodes pairwise sequence similarities of sliding windows from human and mouse syntenic loci, and computation of $G$ can be significantly optimized by minimizing redundant computations in $Eq.1$ coming from overlaps with neighboring sliding windows. Recall that $S_{H,i}$ and $S_{M,j}$ each represent $i^{th}$ and $j^{th}$ sliding windows of $S_H$ and $S_M$. The algorithm requires that $w$ (sliding window width) is divisible by $s$ (slide step size). Then, $S_{H,i}$ and $S_{M,j}$ can each be represented as a concatenated ($\frown$) series of contiguous $s-$mers, repeating $w/s$ times **(Figure 2)**.

[6] Ghandi, Mahmoud, et al. "Enhanced regulatory sequence prediction using gapped k-mer features." PLoS computational biology 10.7 (2014): e1003711.

[7] Throughout this document, all sequence and matrix indices will start from 1.

[8] S(i,n) denotes length-n subsequence of S starting from $i^{th}$ position of S. i.e., S[i+j]=S(i,n)[1+j] for $j \in [0, ..., n-1]$

That is:

$$S_{H,i} = S_H(1+(i-1)\cdot s, s)^\frown S_H(1+i\cdot s, s)^\frown S_H(1+(i+1)\cdot s, s)^\frown \ldots {}^\frown S_H(1+(i+w/s-2)\cdot s, s)$$
$$S_{M,j} = S_M(1+(j-1)\cdot s, s)^\frown S_M(1+j\cdot s, s)^\frown S_M(1+(j+1)\cdot s, s)^\frown \ldots {}^\frown S_M(1+(j+w/s-2)\cdot s, s)$$

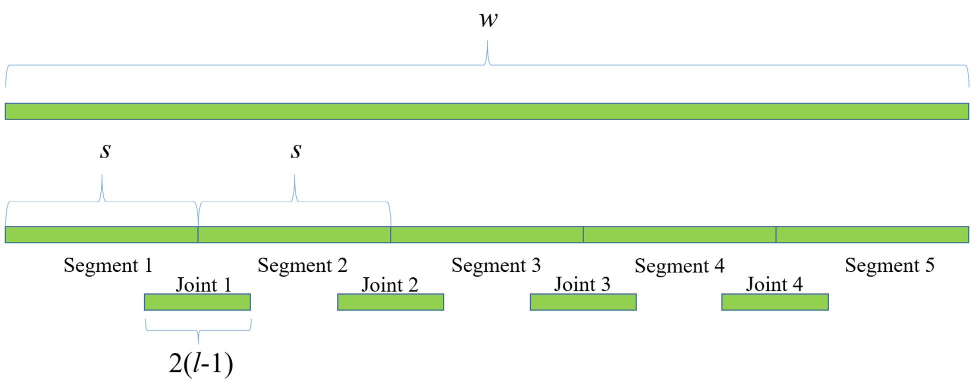


Figure 2: Gapped-kmer composition of a genomic window is equal to the sum of gapped-kmer composition of all its segments and its joint components. Diagram given for $w = 100$ and $s = 20$

Before considering gkm-composition of the above two representations, let's consider the following observation. Suppose $S = S_A^\frown S_B$, where $w_A = |S_A|, w_B = |S_B|, s \geq l$. Then, $g(S) \geq g(S_A) + g(S_B)$ since $S$ contains not only subsequences of A and B but also subsequences that are newly introduced by joining A and B together. Let's call such subsequence of $S$ as $joint(S_A, S_B) = S(w_A - l + 1, 2(l-1))$. Then

$$\begin{aligned} g(S) &= g(S_A^\frown S_B) \\ &= g(S_A) + g(S_B) + g(joint(S_A, S_B)) \end{aligned}$$

Using the same idea, then

$$\begin{aligned} g(S_{H,i}) &= g(S_H(1+(i-1)\cdot s, s)^\frown S_H(1+i\cdot s, s)^\frown \ldots {}^\frown S_H(1+(i+w/s-2)\cdot s, s) \\ &= \sum_{p=1}^{w/s} g(S_H(1+(i+p-2)\cdot s, s)) \\ &+ \sum_{q=1}^{w/s-1} g(joint(S_H(1+(i+q-2)\cdot s, s), S_H(1+(i+q-1)\cdot s, s))), \end{aligned}$$

where the first series computes the total gapped-kmer composition at individual segments being concatenated and the second series computes the total composition at the joints. Naming the terms in the first series as segments and those in the second series as joints for simplicity, we get the following representations for the gapped-kmer compositions of the human and mouse sliding windows.

$$g(S_{H,i}) = \sum_{p=1}^{w/s} g(p^{th}\text{segment of } S_{H,i}) + \sum_{q=1}^{w/s-1} g(q^{th}\text{joint of } S_{H,i}) \qquad \text{(Eq.3)}$$

$$g(S_{M,j}) = \sum_{p=1}^{w/s} g(p^{th}\text{segment of } S_{M,j}) + \sum_{q=1}^{w/s-1} g(q^{th}\text{joint of } S_{M,j}) \qquad \text{(Eq.4)}$$

Dot product of Eq.3 and Eq.4 is then

$$
\begin{aligned}
g(S_{H,i}) \cdot g(S_{M,j}) = & (\sum_{p_1=1}^{w/s} g(p_1^{th}\text{segment of } S_{H,i}) + \sum_{q_1=1}^{w/s-1} g(q_1^{th}\text{joint of } S_{H,i})) \\
& \cdot (\sum_{p_2=1}^{w/s} g(p_2^{th}\text{segment of } S_{M,j}) + \sum_{q_2=1}^{w/s-1} g(q_2^{th}\text{joint of } S_{M,j})) \\
= & (\sum_{p_1=1}^{w/s} \sum_{p_2=1}^{w/s} g(p_1^{th}\text{segment of } S_{H,i}) \cdot g(p_2^{th}\text{segment of } S_{M,j})) \\
& + (\sum_{p_1=1}^{w/s} \sum_{q_2=1}^{w/s-1} g(p_1^{th}\text{segment of } S_{H,i}) \cdot g(q_2^{th}\text{joint of } S_{M,j})) \\
& + (\sum_{q_1=1}^{w/s-1} \sum_{p_2=1}^{w/s} g(q_1^{th}\text{joint of } S_{H,i}) \cdot g(p_2^{th}\text{segment of } S_{M,j})) \\
& + (\sum_{q_1=1}^{w/s-1} \sum_{q_2=1}^{w/s-1} g(q_1^{th}\text{joint of } S_{H,i}) \cdot g(q_2^{th}\text{joint of } S_{M,j})), \qquad \text{(Eq.5)}
\end{aligned}
$$

where each of the dot products of gapped-kmer vector is computed using Eq.2. The dot products in Eq.5 can then be partitioned into four subsets (according to how they grouped into the four double series in Eq.5) and be organized into a 3D hypermatrix.

$$
\begin{aligned}
& P^{ij} \in \mathbb{N}^{(w/s)\times(w/s)\times 4} \\
P^{ij}[q_1, q_2, 1] &= g(q_1^{th}\text{segment of } S_{H,i}) \cdot g(q_2^{th}\text{segment of } S_{M,j}) \\
P^{ij}[q_1, q_2, 2] &= g(q_1^{th}\text{segment of } S_{H,i}) \cdot g(q_2^{th}\text{joint of } S_{M,j}) \\
P^{ij}[q_1, q_2, 3] &= g(q_1^{th}\text{joint of } S_{H,i}) \cdot g(q_2^{th}\text{segment of } S_{M,j}) \\
P^{ij}[q_1, q_2, 4] &= g(q_1^{th}\text{joint of } S_{H,i}) \cdot g(q_2^{th}\text{joint of } S_{M,j})
\end{aligned}
$$

Then, Eq.5 is equivalent to the equation below (Eq.6).

$$
g(S_{H,i}) \cdot g(S_{M,j}) = \sum_{q_1=1}^{w/s} \sum_{q_2=1}^{w/s} \sum_{k=1}^{4} P^{ij}[q_1, q_2, k] \cdot \mathbb{1}(q_1, q_2, k), \qquad \text{(Eq.6)}
$$

where $\mathbb{1}(q_1, q_2, k)$ is a boolean function enforcing the index ranges of the double series in Eq.5.

$$
\mathbb{1}(q_1, q_2, k) = \begin{cases} 1 & , k = 1 \\ 1 \text{ iff } q_2 < w/s & , k = 2 \\ 1 \text{ iff } q_1 < w/s & , k = 3 \\ 1 \text{ iff } q_1, q_2 < w/s & , k = 4 \end{cases}
$$

Note that the entries of $P^{ij}$ are redundantly shared with $P^{xy}$ from neighboring pairs of sliding windows (e.g., $P^{(i-1)j}$,$P^{i(j-1)}$,$P^{(i-1)(j-1)}$, etc). Define matrix $P$, which has every $P^{ij}$ as its submatrices and non-redundantly contains all the entries of $P^{ij}$ for all $i^{th}$ and $j^{th}$ sliding windows.

$$
\begin{aligned}
& P \in \mathbb{N}^{(|S_H|/s)\times(|S_M|/s)\times 4} \\
& P[(i-1) + q_1, (j-1) + q_2, :] = P^{i,j}[q_1, q_2, :] \text{ , where } 1 \leq q_1, q_2 \leq w/s,
\end{aligned}
$$

which leads to:

$$P[(i-1)+q_1,(j-1)+q_2,1] = g(q_1^{th}\text{segment of } S_{H,i})\cdot g(q_2^{th}\text{segment of } S_{M,j}))$$
$$P[(i-1)+q_1,(j-1)+q_2,2] = g(q_1^{th}\text{segment of } S_{H,i})\cdot g(q_2^{th}\text{joint of } S_{M,j})$$
$$P[(i-1)+q_1,(j-1)+q_2,3] = g(q_1^{th}\text{joint of } S_{H,i})\cdot g(q_2^{th}\text{segment of } S_{M,j}))$$
$$P[(i-1)+q_1,(j-1)+q_2,4] = g(q_1^{th}\text{joint of } S_{H,i})\cdot g(q_2^{th}\text{joint of } S_{M,j}))$$

Finally, Eq.6 is equivalently and more compactly expressed as:

$$g(S_{H,i})\cdot g(S_{M,j}) = \sum_{q_1=1}^{w/s}\sum_{q_2=1}^{w/s}\sum_{k=1}^{4} P[(i+q_1-1),(j+q_2-1),k]\cdot \mathbb{1}(q_1,q_2,k) \qquad \text{(Eq.7)}$$

The matrix $P$ contains all necessary and sufficient information for computing $G_{S_H,S_M}[i,j]$ for all pairs of $i^{th}$ and $j^{th}$ sliding windows of $S_H$ and $S_M$. It contains gapped-kmer similarities of all pairs of atomic fragments of $S_H$ and $S_M$ that can be combinatorially summed to compute each elements of $G_{S_H,S_M}$. Each element in $P$ is non-redundantly computed and stored. $P$ is initialized to be a matrix of -1, and its elements are replaced with a non-negative number while computing $G$. For example, P[1,2,1] is needed both for computing $g(S_{H,1},S_{M,1})$ (via $i,j,q_1,q_2 = 1,1,1,2$) and for computing $g(S_{H,1},S_{M,2})$ (via $i,j,q_1,q_2 = 1,2,1,1$), but it is computed once while computing $g(S_{H,1})\cdot g(S_{M,1})$. All elements of P are computed once and stored during the course of computing G, and they are accessed when they are needed for computing other elements of G. This optimization makes the time-complexity of gkm-align nearly independent of the sliding window width ($w$).

Hence, the results from **B.1** and **B.2** give gkm-align algorithm the time-complexity independent of both $l$ (gapped-kmer width) and $w$ (window width). It is dependent on the sizes of the genomes and sizes of the human-mouse syntenic blocks (**Fig.1D**, **section D**).

### C Finding the optimal alignment path of maximum gapped-kmer similarity using matrix $G$

After computing matrix $G$, as described in **section B**, we can now compute the optimal collinear alignment path spanning the human and mouse genomic loci at the resolution of sliding windows. Resulting alignment path is a collinear sequence of pairs of human and mouse genomic windows that most likely contain conserved DNA elements inherited from the human-mouse common ancestor.

Provided with matrix $G \in [0,1]^{m \times n}$ that contains pairwise sequence-similarities of every pair of sliding windows in $S_H$ and $S_M$, we define a collinear path of sliding windows as a sequence of matrix indices [9] $P = ((i_1, j_1), (i_2, j_2) \ldots, (i_{|P|}, j_{|P|}))$, such that

$$
\begin{aligned}
&(i_1, j_1) = (1, 1) && (1)\\
&(i_{|P|}, j_{|P|}) = (m, n) && (2)\\
&(i_{k+1}, j_{k+1}) - (i_k, j_k) \in \{(1,1), (1,0), (0,1)\} && (3)\\
&p \geq q \implies i_p \geq i_q \text{ and } j_p \geq j_q, && (4)
\end{aligned}
$$

where $(i_{k+1}, j_{k+1}) - (i_k, j_k) \in \{(1,0), (0,1)\}$ indicates insertion or deletion events in $S_H$ or $S_M$ relative to each other. To find the most likely path that describes the evolutionary history between $S_H$ and $S_M$, we use a variant of the Smith-Waterman algorithm. To find the optimal path $P$, we score all possible paths satisfying (1-4) as follows:

$$f(P) = \sum_{q=1}^{|P|} G(P(i)) - \alpha \cdot I,$$

where $I$ is the number of indel events in $P$.

$$I = |\{k \in 1 \ldots |P| - 1 : (i_{k+1}, j_{k+1}) - (i_k, j_k) \in \{(1,0), (0,1)\}\}|$$

It is difficult to choose a stable value for $\alpha$ since it depends on the parameters used for computing the sequence similarities, such as the width of the sliding windows ($w$) and $l$ and $k$ for the gapped-kmer definition. To choose $\alpha$ independent of $(w, l, k)$, we Z-transform matrix $G$ so that

$$f(P) = \sum_{q=1}^{|P|} \frac{G(P(q)) - \mu_G}{\sigma_G} - \alpha \cdot I_q \qquad \text{(Eq.8)}$$

Z-transforming also has an additional advantage of normalizing sequence similarities relative to local background sequence similarities, which are variable by loci, for example, by GC contents. Globally optimal $P$ for a given $G$ is then efficiently identified using dynamic programming.

$$
\begin{aligned}
\text{optimal alignment path spanning } S_H \text{ and } S_M &= argmax_P f(P)\\
&= argmax_P \sum_{q=1}^{|P|} \frac{G(P(q)) - \mu_G}{\sigma_G} - \alpha \cdot I_q\\
&= argmax_P \sum_{q=1}^{|P|} \frac{G(P(q)) - \mu_G}{\sigma_G} \text{ , if } \alpha{=}0
\end{aligned}
$$

[9] The matrix indices are eventually converted to human-mouse genomic coordinates for gkm-align output.

### D Whole-genome extension of gkm-align

**Section A** through **C** explained how local gkm-alignment is performed (**Figure 1A-C**) given that the human-mouse syntenic blocks of interest are specified (visualized as the rectangles in **Figure 1D**). Here, we discuss how gkm-align is extended genome-wide. The essence of whole-genome extension is the identification of human-mouse syntenic blocks. Most conventional genome-alignment methods [10,11,12] identify syntenic blocks by (1) identifying short sequence matches between human and mouse genomes (2) and chaining the matches based on their genomic coordinates. Gkm-align adopts this idea.

First, we obtained short sequence matches (*seed hits*) between human and mouse using LASTZ (v1.04.04; options: –notransition –step=1 –gfextend –nochain –nogapped) (visualized by the dots in **Figure 1D**). Then, to focus on conserved gene regulatory circuits, we filtered out seed hits that are not located within any human/mouse syntenic intergenic locus identified using the ENCODE gene ortholog list [13] (**Methods**).

Then, to derive syntenic blocks, we implemented the chaining algorithm described by Zhang et al[14] with slight modifications. The chaining algorithm returns a list of seed hits partitioned into syntenic blocks(e.g., the three dot colors in **Figure 1D**), and seed hits at the two extremes of each syntenic blocks can be used to define genomic ranges of human/mouse syntenic loci. But there can be abnormally large syntenic blocks, and computing matrix $G$ of these blocks can be computationally infeasible. We further break down the syntenic blocks by clustering the seed hits by their genomic coordinates within the identified syntenic blocks (number of centroids determined by targeted average syntenic block size), and the seed hits nearest to the centroids are then used to define smaller syntenic blocks (the rectangles in **Figure 1D**). Gkm-align then computes matrix $G$ for each of the smaller syntenic blocks for local alignment. Gkm-align software provides an option to compute matrix $G$s in parallel with multithreading.

### E Enhanced discovery of conserved enhancers by incorporating cell-specific regulatory vocabularies

We can further enhance discovery of orthologous enhancer pairs using sequence-based machine learning models. Gkm-SVM models can distinguish enhancers from random genomic background, and their prediction values for each human and mouse elements can be weighted into sequence similarity matrix G to enhance alignment.

Recall that sequence similarity between $i^{th}$ and $j^{th}$ slidindg windows of sequences $S_H$ and $S_M$ (denoted as $S_{H,i}, S_{M,j}$) is computed as

$$G_{S_H,S_M}[i,j] = \frac{g(S_{H,i}) \cdot g(S_{M,j})}{|g(S_{H,i})||g(S_{M,j})|}$$

Then, gkm-SVM weighted sequence similarity is computed as

$$G_{S_H,S_M}[i,j] \leftarrow (G_{S_H,S_M}[i,j])^{1-c}(P_M(S_{H,i}) \cdot P_H(S_{M,j})))^c,$$

where $P_M$ and $P_H$ are gkm-SVM enhancer prediction functions ($P : \mathit{sequence} \rightarrow [0,1]$) trained on mouse (M) and human (H) enhancers. Higher $c \in [0,1]$ indicates higher influence of gkm-SVM predictions on alignment.

Section **E.1** and **E.2** describe how $P_H()$ and $P_M()$ are computed. Section **E.3** describes how a similar idea using gkm-SVM is used to mask repetitive DNA elements prior to matrix $G$ computation.

[10]Delcher, Arthur L., et al. "Alignment of whole genomes." Nucleic acids research 27.11 (1999): 2369-2376.
[11]Kent, W. James, et al. "Evolution's cauldron: duplication, deletion, and rearrangement in the mouse and human genomes." Proceedings of the National Academy of Sciences 100.20 (2003): 11484-11489.
[12]Harris, Robert S. Improved pairwise alignment of genomic DNA. The Pennsylvania State University, 2007.
[13]Yue, Feng, et al. "A comparative encyclopedia of DNA elements in the mouse genome." Nature 515.7527 (2014): 355-364.
[14]Zhang, Zheng, et al. "Chaining multiple-alignment blocks." Journal of Computational Biology 1.3 (1994): 217-226.

#### E.1 Using gkm-SVM kmer-weight vectors for enhancer prediction

Training gkm-SVM on enhancers generates support vector sequences ($SV$) and corresponding support vector weights ($\alpha$), which can be used to make enhancer predictions on an arbitrary DNA sequence $X$.

$$\begin{aligned} pred(X) &= \sum_{S \in SV} \alpha_S < g(X), g(S) > \\ &= \sum_{S \in SV} \alpha_S \frac{g(X) \cdot g(S)}{|g(X)||g(S)|} \end{aligned}$$

Further, by computing gkm-SVM predictions on every kmers, we can quantify regulatory importance of each kmers (default kmer-width = 11; e.g., CCATGGCAACC: top RFX binding motif for brain enhancers). Gkm-SVM weight of a kmer sequence $k_{mer}$ is computed as:

$$w_{k_{mer}} = pred(k_{mer}) = \sum_{S \in SV} \alpha_S < g(k_{mer}), g(S) > \qquad \text{(Eq.7)}$$

Pre-computed gkm-SVM kmer weights ($w_{kmer}$) can be used to efficiently compute gkm-SVM prediction score for sequence $X$ by its kmer weight sum.

$$\begin{aligned} pred(X) &= \sum_{S \in SV} \alpha_S < g(X), g(S) > \\ &= \sum_{S \in SV} \alpha_S \frac{g(X) \cdot g(S)}{|g(X)||g(S)|} \\ &= \sum_{S \in SV} \alpha_S \frac{(\sum_{k_{mer} \in X} g(k_{mer})) \cdot g(S)}{|g(X)||g(S)|} \\ &= \sum_{k_{mer} \in X} \sum_{S \in SV} \alpha_S \frac{g(k_{mer}) \cdot g(S)}{|g(X)||g(S)|} \\ &= \frac{|g(k_{mer})|}{|g(X)|} \sum_{k_{mer} \in X} \sum_{S \in SV} \alpha_S \frac{g(k_{mer}) \cdot g(S)}{|g(k_{mer})||g(S)|} \\ &= \frac{|g(k_{mer})|}{|g(X)|} \sum_{k_{mer} \in X} \sum_{S \in SV} \alpha_S < g(k_{mer}), g(S)) > && \text{(use Eq.2 for } |g(k_{mer})|, |g(X)|) \\ &= \frac{\sqrt{\binom{l}{k}}}{\sqrt{(|X| - l + 1)^2 \binom{l}{k}}} \sum_{k_{mer} \in X} \sum_{S \in SV} \alpha_S < g(k_{mer}), g(S)) > && \text{(use Eq.7 for } w_{k_{mer}};\ |X| = w) \\ &= \frac{1}{w - l + 1} \sum_{k_{mer} \in X} w_{k_{mer}} && (w,l \text{ are constants in gkm-align}) \\ &\propto \sum_{k_{mer} \in X} w_{k_{mer}} && \text{(Eq.8)} \end{aligned}$$

#### E.2 Probabilistic interpretation and normalization of gkm-SVM prediction

One limitation of directly incorporating gkm-SVM predictions to gkm-align is that these values are not bounded, limiting interpretable application for genome alignment. Instead, we transform the prediction values to range between 0 and 1 by computing the posterior probability that a sequence is an enhancer. This is done by computing gkm-SVM score distributions for enhancers and non-enhancers (i.e., positive and negative training sets used for gkm-SVM enhancer training; prediction scores computed with 5-fold cross validation)

$$\begin{aligned} \mu_{(+)} &= E(pred(X_{pos})), \sigma^2_{(+)} = Var(pred(X_{pos})) \\ \mu_{(-)} &= E(pred(X_{neg})), \sigma^2_{(-)} = Var(pred(X_{neg})) \end{aligned}$$

Assuming that priors for finding positive and negative sequences are equal, we compute posterior gkm-SVM prediction score of a given sequence X with

$$P_{gkm}(X) = P(X \in X_{pos} | pred(X)) = \frac{P_{(+)}}{P_{(+)} + P_{(-)}} \in [0, 1], \qquad \text{(Eq.9)}$$

where

$$P_{(+)} = \frac{1}{\sqrt{2\pi\sigma^2_{(+)}}} exp(\frac{pred(X) - \mu_{(+)}}{2\sigma^2_{(+)}})$$

$$P_{(-)} = \frac{1}{\sqrt{2\pi\sigma^2_{(-)}}} exp(\frac{pred(X) - \mu_{(-)}}{2\sigma^2_{(-)}})$$

$pred(X)$ is computed using Eq.8, summing up kmer-weights of every kmer contained in $X$.

##### E.3 Detecting and masking repetitive elements using gkm-SVM

Without masking repetitive DNA elements, Matrix $G$ is riddled with sequence similarities coming from high-entropy repetitive elements that are prevalent across the human and mouse genomes (mostly simple low-complexity repeats; **Supp Fig.8**). Hence, a method to underweight sequence similarities coming from repetitive elements is crucial for reliable genome alignment.

One intuitive solution is to use principal component analysis on the rows and columns of $G$ to automatically detect elements in $S_H$ that are similar to many elements in $S_M$ and vise versa. However, such method is computationally too inefficient to apply genome-wide. Instead, we can again use sequence-based machine learning methods (e.g. gkm-SVM) to learn patterns of DNA sequences that are highly populated in the human and mouse genomes, and use the model to detect and filter out repetitive elements in linear time. We first train gkm-SVM, separately for human and mouse, on randomly sampled genomic windows (300 base pairs wide) outside regulatory regions (e.g., union of DHS across all ENCODE experiments) against randomly generated sequences such that each base pair has equal probabilities of being A,C,G, or T. This results in a kmer-weight vector, where kmers with high weight are significantly enriched in the genome more than expected by chance (**Supp Fig.8A**; e.g., ATATATATATA).

In order to detect base pairs that highly contribute to repetitiveness observed in matrix $G$, we compute repeat scores of each base pairs in the genome by summing up kmer weights of kmers that overlap with the base pair. For $i^{th}$ base pair in sequence $S$, its repeat score is computed as

$$r(S[i]) = \sum_{j=i-k+1}^{i} w_{S(j,k)},$$

where k = kmer size. Using this function, we can compute repeat scores for each base pairs across the entire genome, and mask base pairs that receive repeat scores above a certain threshold, determined by the percentage of the genomic base pairs that we target to mask. For gkm-align, we used a repeat score threshold that would mask approximately 10 percent of the human and mouse genomes. Base pairs that surpass the estimated threshold are masked by replacement with random characters. They are replaced with ASCII characters excluding (A, C, G, T) in order to nearly eliminate the chance that the newly introduced characters contribute any artificial sequence similarities. After repeat masking, we compute matrix $G$ as described in **section B**, compute optimal alignment paths using $G$ (**section C**), and repeat this process across every human-mouse syntenic blocks (**section D**) for whole-genome alignment.
